# The dynamic and diverse nature of parenchyma cells in the Arabidopsis root during secondary growth

**DOI:** 10.1101/2024.07.18.604073

**Authors:** Munan Lyu, Hiroyuki Iida, Thomas Eekhout, Meeri Mäkelä, Sampo Muranen, Lingling Ye, Anne Vatén, Brecht Wybouw, Xin Wang, Bert De Rybel, Ari Pekka Mähönen

**Author notes:** These authors contributed equally (Munan Lyu, Hiroyuki Iida).

## Abstract

During the process of secondary growth, the vascular cambium produces the conductive xylem and phloem cells, while the phellogen (cork cambium) deposit phellem (cork) as the outermost protective barrier. Although most of the secondary tissues is made up by parenchyma cells which are also produced by both cambia, their diversity and function are poorly understood. Here we combined single-cell RNA sequencing analysis with lineage tracing to recreate developmental trajectories of the cell types in the *Arabidopsis* root undergoing secondary growth. By analysing 93 reporter lines, we were able to identify 20 different cell types or cell states, many of which have not been described before. We additionally observed distinct transcriptome signatures of parenchyma cells depending on their maturation state and proximity to the conductive cell types. Our data shows that both xylem and phloem parenchyma tissues are required for normal formation of conductive tissue cell types. Furthermore, we showed that mature phloem parenchyma gradually obtains periderm identity, and this transition can be accelerated by jasmonate or wounding. Thus, our study reveals the remarkable dynamic and diverse nature of parenchyma cells during secondary growth.

## Main

Plants undergo primary growth for longitudinal elongation. Additionally, many seed plants exhibit secondary (i.e. radial) growth in their mature stems and roots orchestrated by the lateral meristems, the vascular cambium and the phellogen (cork cambium). The vascular cambium produces conductive cells, secondary xylem inward and secondary phloem outward, and the phellogen provides a barrier tissue, the phellem (cork) outward. Besides these conductive or barrier cell types, both cambia also generate parenchyma cells, xylem and phloem parenchyma from the vascular cambium and the phelloderm from the phellogen, and these parenchymatic cells occupy the largest part of secondary tissues^1–3^. Parenchyma is generally consisted of thin-walled living cells; there are various types of parenchyma cells and some of their functions are known, for instance, mesophyll cells are essential for photosynthesis^1^. However, because parenchyma cells have received less attention compared to the conductive cells in secondary tissues, their functions are largely unknown. Also, from a morphological point of view, there are very little to no differences among the parenchymatic cells in the root^4^. Although a few reports suggest heterogeneity of xylem parenchyma cells^5^, it has not been proven experimentally and it is unclear even whether parenchyma cells within the same tissue have a heterogenous or homogeneous identity. Altogether, little is known about the parenchyma cell types nor their functions in secondary tissues. Here, we optimised single-cell RNA-sequencing^6–8^ on *Arabidopsis* mature roots to explore the diversity of cell types in root undergoing secondary growth. We validated that our dataset contains all known conductive and parenchymatic cell types in *Arabidopsis* secondary tissues and demonstrate that xylem and phloem parenchyma are composed of diverse cell types and cell states. Through extensive reporter analysis combined with mutant analysis, we also identified that the xylem and phloem parenchyma cells function in supporting conductive tissue formation. Furthermore, lineage tracing analysis suggests that mature phloem parenchyma cells can change their cell identity to replenish a new barrier upon injury. Taken together, our study demonstrates the diverse and dynamic nature of parenchyma cells in *Arabidopsis* secondary tissues.

## Results

### Single-cell transcriptomics profiling revealed a diverse set of cell types in *Arabidopsis* roots during secondary growth

To explore cell type diversity and dynamics during root secondary growth, we produced a single-cell RNA-seq atlas. For this study, we utilized the 30-day-old *Arabidopsis* seedlings that had just started bolting and were characterised by intensive root secondary growth (Fig. 1a). Sections of the first 2 cm below the root-hypocotyl junction were collected for protoplast isolation and subsequent transcriptome profiling (Fig. 1a). After quality control ^7^, 11,760 high-quality cells (with median 3,160 genes and median 14,709 reads per cell) were retained and visualized using uniform manifold approximation (UMAP)^9^ (Fig. 1b). To examine whether the major cell types known to be present in this tissue were captured in our dataset, we initially analysed expression of known tissue-specific genes^10–16^. Based on these markers, all the known cell types were predicted to be present in the dataset, while several clusters remained unidentified (Supplementary Fig. 1a,b). The relative positions of each cell type in the dataset UMAP reflect the real organisation of the secondary root, indicating that it is likely that we captured most of the developmental states and transitions between them (Fig. 1a,b). To validate these predictions and annotate all the remaining clusters, we examined expression of 93 fluorescent reporters: 16 published before^17–24^ and 77 generated for this study (Supplementary Fig. 1c). Many of these reporters indicated the cell states for which no marker had been previously identified and can serve as tissue-specific markers in secondary tissues of *Arabidopsis* roots. A detailed description of the 93 reporter lines used to annotate the clusters is described below and in the Supplementary Fig. 1c-i, Supplementary Notes and Supplementary Table S4. Also, strongly and ubiquitously expressed promoters for overexpression studies in the secondary tissue are presented in the Supplementary Notes and in Supplementary Fig. 1h,i.

**Figure 1.**
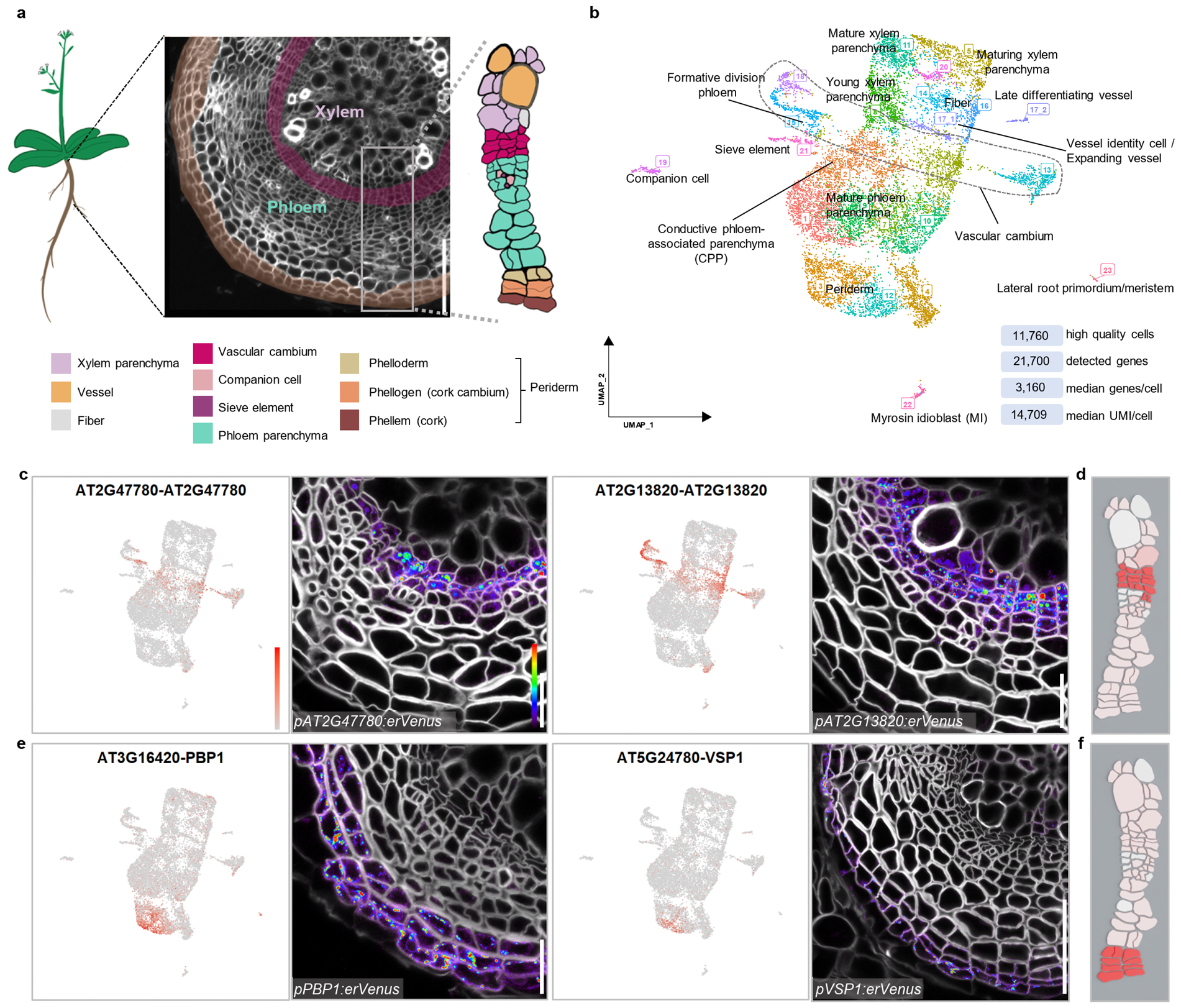
Single-cell transcriptome profiling of the *Arabidopsis* mature root. **a**, Illustration of the tissue utilized for the single-cell RNA sequencing. The illustrative image is a cross section of 30-day-old Arabidopsis mature root, **b**, Visualization of 23 cell clusters using Uniform Manifold Approximation and Projection (UMAP), with identity annotations validated by reporter analysis. Each dot represents an individual cell, with colors representing different clusters. A dotted circle highlights the vascular cambial cells. The dataset quality information is displayed in the bottom right corner, **c**, UMAP plots of genes (AT2G4 7780 and AT2G13820) specifically detected in vascular cambium clusters and confocal cross-sections of their promoter-reporter lines, **d**, Expression of AT2G13820 in Celia model, **e**, UMAP plots of genes (PBP1 and VSP1) highly detected in periderm clusters and confocal cross-sections of their promoter-reporter lines, **f,** Expression of VSP1 in Celia model. In cross sections of **a, c, e,** cell walls were stained with SR2200. In c and e, for each gene, the UMAP plot and cross section of 16-day-old roots was shown in the left and right, respectively. Relative expression levels of genes in UMAP and Venus-YFP signals in cross sections were shown according to the colour map in **c**. Scale bars, 100 pm (**a**), 20 pm **(c, e).**

Vascular cambium and phellogen (cork cambium) are secondary meristems which orchestrate radial growth^1^. Since some key vascular cambium regulators are expressed in the periderm (that is, phellem, phellogen and phelloderm) as well, these two cambia share core developmental regulators^2^. However, the different function between these two cambia also indicates that they have cambium-specific regulators. Our single-cell RNA-seq data captured numerous cells of both cambia and successfully identified genes unique to one cambium, as well as those expressed in both (Fig. 1c-f, Supplementary Fig. 1d,e, Supplementary Fig. 2 and Supplementary Notes). Taken together, we generated and validated a transcriptome atlas of the *Arabidopsis* root undergoing secondary growth; allowing to reveal shared and unique regulatory mechanisms driving vascular cambium and/or phellogen activity. This atlas is available in an online tool (https://www.single-cell.be/plant) together with model based on Cella^25^ (Fig. 1d,f).

### The xylem parenchyma cell pool consists of at least three distinct developmental states

Next, we analysed different cell types in the xylem domain by combining bioinformatic and reporter analysis. We identified xylem fibers and xylem vessels at various stages of development in the dataset. Subcluster 17_1 represents cells that have recently obtained vessel identity and initiated cell expansion, and from there cells progress to subcluster 17_2 to finalize terminal differentiation (Supplementary Notes, Supplementary Fig. 3 and Supplementary Fig. 4a).

Given that xylem parenchyma is less comprehensively understood compared to the morphologically distinct xylem cell types, we conducted a deeper analysis on xylem parenchyma. We found that xylem parenchyma cells were classified in three clusters (8, 11 and 5) containing different transcriptional information. The transcriptional reporter lines of genes predominantly expressed in cluster 8 (*ROTUNDIFOLIA LIKE 6* (*RTFL6*), *MIZU-KUSSEI 1* (*MIZ1*), and *AT1G03620*) showed high fluorescence signals in young xylem parenchyma cells adjacent to the meristematic zone and near the expanding vessels (Fig. 2a and Supplementary Fig. 4b). Reporter expression driven by the promoters of cluster 11-enriched genes (*AT5G07080* and *AT1G11925*) were preferentially detected in the most mature xylem parenchyma cells near the primary xylem axis both in 16-day-old and in 30-day-old roots (Fig. 2c and Supplementary Fig. 4c,d). The cluster 5-enriched genes (*AT1G48750*, *UDP-GLUCOSYL TRANSFERASE 72D1* (*UGT72D1*) and *CASPARIAN STRIP INTEGRITY FACTOR 2* (*CIF2*)) were expressed in the parenchyma between the mature and young xylem parenchyma cells, which we termed maturing xylem parenchyma (Fig. 2b and Supplementary Fig. 4e). These results thus suggest that the xylem parenchyma is not a homogenous tissue but is rather composed of cells with different maturation states. To further characterize the different states, we performed gene ontology (GO) comparison of xylem parenchyma clusters. Cluster 8 showed specific response to salt stress, auxin transport and cluster 11 to hypoxia, additionally both clusters were enriched by genes, for example, related to response to fungus (Supplementary Fig. 4f). Cluster 5 DEGs were involved in phenylpropanoid biosynthesis and metabolic pathways, that are required for processes such as lignin biosynthesis and defence responses^26^(Supplementary Fig. 4f). As such, the subsequent maturation states of the xylem parenchyma appear to be correlated with different functions including stress responses and vessel lignification. Supporting this, it has been shown that neighbouring xylem parenchyma participate in vessel lignification^27^. To furhter investigate the function of xylem parenchyma cells, we examined a series of T-DNA insertion mutants whose corresponding genes are expressed preferentially in any of the states of xylem parenchyma maturation (Fig. 2d and Supplementary Fig. 4g). Even though the expression of these genes was low in the vessel cluster, nearly half of examined single mutants (four out of nine mutants) showed a significant reduction in secondary vessel formation (Fig. 2e,f and Supplementary Fig. 4h,i). Secondary growth in these mutants tended to be reduced, however, even when considering their growth retardation, secondary vessel formation was proportionally significantly decreased in *at4g30460* and *apk1a* loss-of-function mutants (Fig. 2e and Supplementary Fig. 4h,i). Taken together, our results indicate that xylem parenchyma cells are diverse, and one of the functions of xylem parenchyma is to promote vessel formation during the secondary growth.

**Figure 2.**
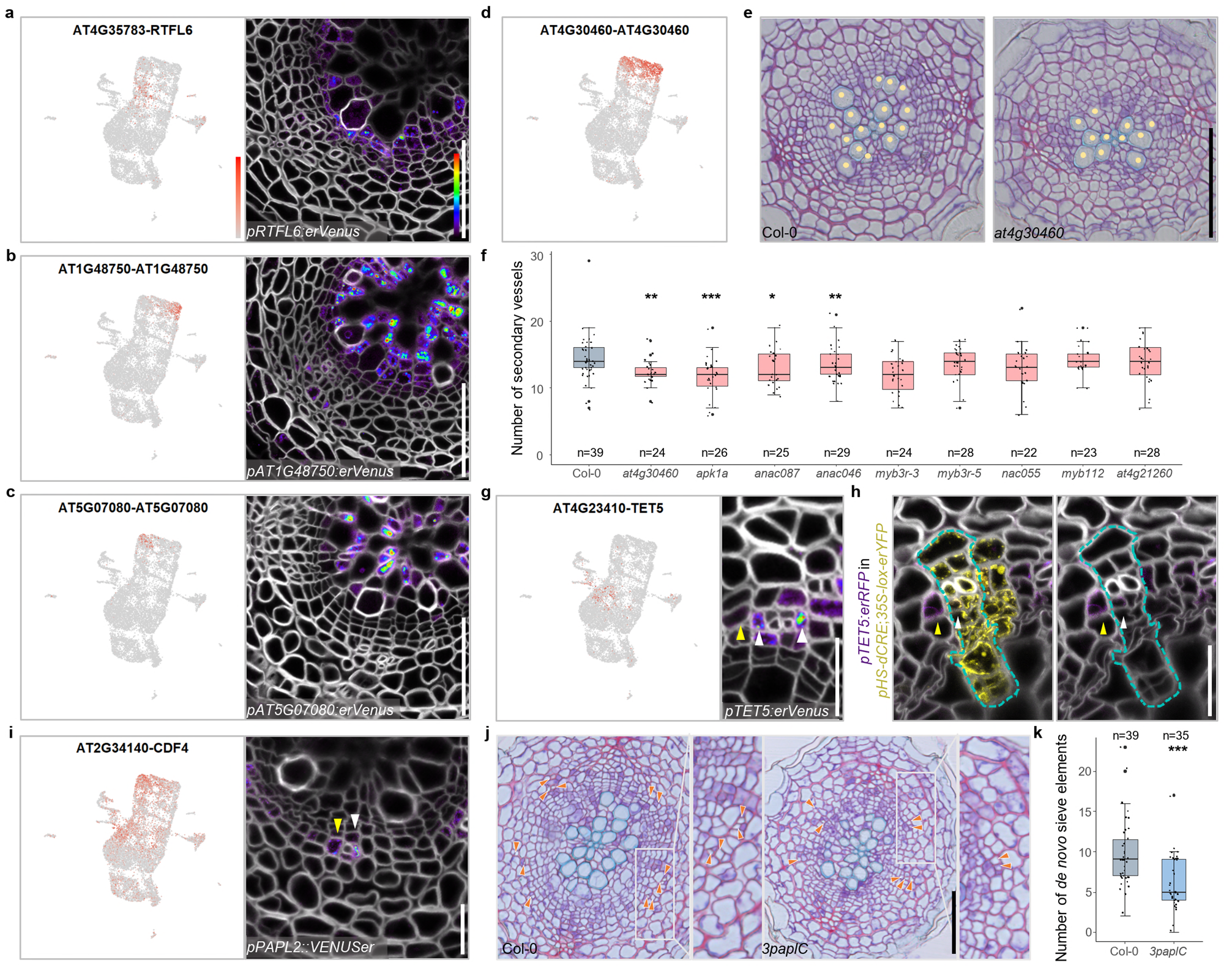
Diversity of parenchyma cells and their functions in conductive tissue development. **a,b,c,** UMAP plots of RTFL6 **(a),** AT1G48750 **(b)** and AT5G07080 **(c)** highly detected in young, maturing, and mature xylem parenchyma cluster, respectively, and confocal cross-sections of their promoter-reporter lines, **d,** UMAP plot of AT4G30460 highly expressed in xylem parenchyma clusters, **e,** Bright-field cross-sections of 14-day-old wildtype and at4g30460 single mutant roots. Yellow dots indicate secondary vessels, **f**, Quantification of secondary vessels number in 14-day-old wildtype and mutant seedlings, **g**, UMAP plot of TET5 highly detected in CPP cluster and confocal cross-section of its promoter reporter line, **h**, Confoc al cross-section of 16-day-old pTET5:erRFP in pHS-dCRE,35S-lox-erYFP root 6 days after clone induction. Left; merged image, right; image with RFP and cell wall channels. A sector from a single clone was encircled by dotted lines. Yellow; Venus-YFP signals, **i**, UMAP plot of CDF4/PAPL1 and confocal cross-sections of its promoter reporter line. In **g-i**, white and yellow arrowheads indicate CPP cells originated from formative division and recruited from neighboring cell lineages, respectively, **j**, Bright-field cross sections of 16-day-old wildtype and 3paplC mutant roots. Orange arrowheads point at de novo sieve elements, **k**, Quantification of de novo sieve elements number in 16-day-old wildtype and 3paplC mutant seedlings. In a-c and **g, i**, for each gene, the UMAP plot and cross section of 16-day-old roots was shown in the left and right, respectively. In cross sections of **a-c** and **g-i**, cell walls were stained with SR2200. In **f** and **k**, the boxes in the box-and-whisker plots represent median values and interquartile range, the whiskers indicate the total range. The black dots indicate measurements from individual roots. Shapiro-Wilk normality test followed by a two-tailed Wilcoxon test was utilized for statistic test between Col-0 and mutants. *p<0.05, **p<0.01, ***p<0.001. The experiment was repeated twice **(f)** or four times **(k)**. n indicate the number of examined roots. Relative expression levels of genes in UMAP and Venus-YFP signals in **a-d**, **g,** and **i** or RFP signals in h in cross sections were shown according to the colour map in a. Scale bars, 50 pm **(e, j),** 20 pm **(a-c, g-i).**

### A subset of phloem parenchyma cells controls conductive phloem formation

Next, we investigated the cell types in phloem domain by integrating bioinformatic and reporter analysis with lineage tracing. Similar to the primary phloem differentiation process in the root tip^17,18^, the formative division in phloem identity cells (subgroup of cluster 15) give rise to sieve elements (cluster 21) and companion cells (cluster 19) (Supplementary Notes and Extended Fig. 5a-c).

Since most secondary phloem tissue consists of parenchyma cells (Fig. 1a) and we know very little about this cell type, we carried out detailed analysis of the phloem parenchyma cell clusters. During the validation process, we noticed that the promoter activities of the phloem-side cambium genes, *DNA BINDING WITH ONE FINGER 2.4* (*DOF2*.4) / *PHLOEM EARLY DOF 1* (*PEAR1*) ^11,17^ and *AT1G12080*, were also highly detected in the parenchyma cells adjacent to the sieve elements and companion cells (Supplementary Fig. 5d). This expression implies the unique identity of phloem parenchyma associated with conductive phloem cells. This type of expression pattern seems to be associated with cluster 2 since the reporters of cluster 2-enriched genes (*TETRASPANIN5* (*TET5*), *AT3G16330* and *WRKY DNA-BINDING PROTEIN 63* (*WRKY63*)) showed preferential expression in those parenchyma cells (Fig. 2g and Supplementary Fig. 5e). We named these cells in cluster 2 as <u>C</u>onductive <u>P</u>hloem-associated <u>P</u>arenchyma (CPP) cells. Next, we studied the ontogeny of these CPP cells and devised two possible mechanisms. First, CPP identity could be established by lineage-based mechanisms by which these cells are co-ordinately formed with the conductive phloem within the same lineage. Alternatively, these cells could be recruited by cell-cell communication-based mechanism from neighbouring conductive cells regardless of their lineage. To examine between these two possibilities, we generated *pTET5:erRFP* reporter line in the background of the cell-lineage tracing line in which the clones were marked with the expression of *YFP* reporter gene upon heat shock (modified from Smetana et al ^11^). Sector analysis was performed six days after *YFP* clones were induced in 10-day-old roots (Fig. 2h). *TET5* signal was detected in CPP cells of both the conductive and nonconductive phloem lineage (Fig. 2h). Thus, both mechanisms coexist; CPP identity can be obtained as a result of formative divisions of conductive cell types as well as recruitment by the neighbouring conductive phloem in a non-cell autonomous manner. Location of the CPP cells reminded us of genes expressed in the cells surrounding the conductive phloem in primary roots such as *PINEAPPLE* (*PAPL*) transcription factors^19^. In our UMAP, three *PAPL* genes showed high expression in cluster 2, and, consistently, fluorescence signals in their reporter lines were detected in CPP cells (Fig. 2i and Supplementary Fig. 5f). While there was a slight but significant reduction in secondary growth in the *papl* triple mutant (Fig. 2j and Supplementary Fig. 5g), a strong reduction in sieve element formation outside of the primary phloem poles region (i.e. *de novo* sieve element formation) was observed (Fig. 2j,k). The ratio of sieve elements versus vasculature diameter reduced significantly in the triple mutant (Supplementary Fig. 5g), implying that the reduction in *de novo* secondary sieve element formation in the *papl* triple mutant is not caused by general growth retardation. These findings suggest that *PAPL* genes are required for the conductive phloem formation. Taken together the findings in this and the previous subsection, disruption of genes expressing in xylem or phloem parenchyma cells resulted in less conductive cell formation, suggesting that xylem and phloem parenchyma generally support conductive cell formation.

### Mature phloem parenchyma is involved in biotic stress responses

As continuous growth pushes the more mature phloem parenchyma cells outwards, these cells become larger and more loosely arranged than the young parenchyma cells near vascular cambium. Based on this morphological characteristic, we named these large parenchyma cells ‘mature phloem parenchyma’ cells, a novel cell type marked by *DEFECTIVE IN INDUCED RESISTANCE 1* (*DIR1*), *COLD-REGULATED 15A* (*COR15A*), and *AT1G62500* expression (Fig. 3a and Supplementary Fig. 6a). Within this region, we identified the existence of a specialized parenchymatic storage cell, myrosin idioblast (MI), which functions as part of the unique glucosinolate-myrosinase defence system in *Brassicaceae* plants against hervivory attacks^28–30^. Majority of validated or potential regulators of MI function or differentiation^31^ showed strong expression in the MI cluster (22 out of 35 genes; Supplementary Fig. 6c). Also, signals of myrosinase (also called thioglucosidase) encoding gene *THIOGLUCOSIDE GLUCOHYDROLASE2* (*TGG2*) ^32^and MI regulator *FAMA*^30,31^ reporters were sparsely detected in individual cells in mature phloem parenchyma region (Fig. 3b and Supplementary Fig. 6b). We utilized Coomassie brilliant blue (CBB) staining to visualize the Mis^33^, and supportively, CBB-stained cells were scattered in the similar area where *TGG2* and *FAMA*-positive cells were found (Fig. 3c). To our knowledge, this is the first report demonstrating that MI exists also in the mature root. This also highlights the power of single cell approaches to reveal the presence of a cell type which has not been reported before in mature root. Interestingly, we often detected promoter activities of *TGG2* and *FAMA*, and CBB staining, in cells next to the sieve element, the position previously occupied by the companion cells (Fig. 3b,c and Supplementary Fig. 6b). From GO enrichment comparison among conductive phloem clusters (cluster 2, 19, 21) and cluster 22, cluster 22 companion cells seem to have a tight connection with MI cells in the context of glucosinolate catabolic and sulfur compound metabolic processes (Supplementary Fig. 6d). In summary, although we cannot exclude other origins, our data strongly supports the notion that MI cells mature from companion cells in the phloem lineage.

**Figure 3.**
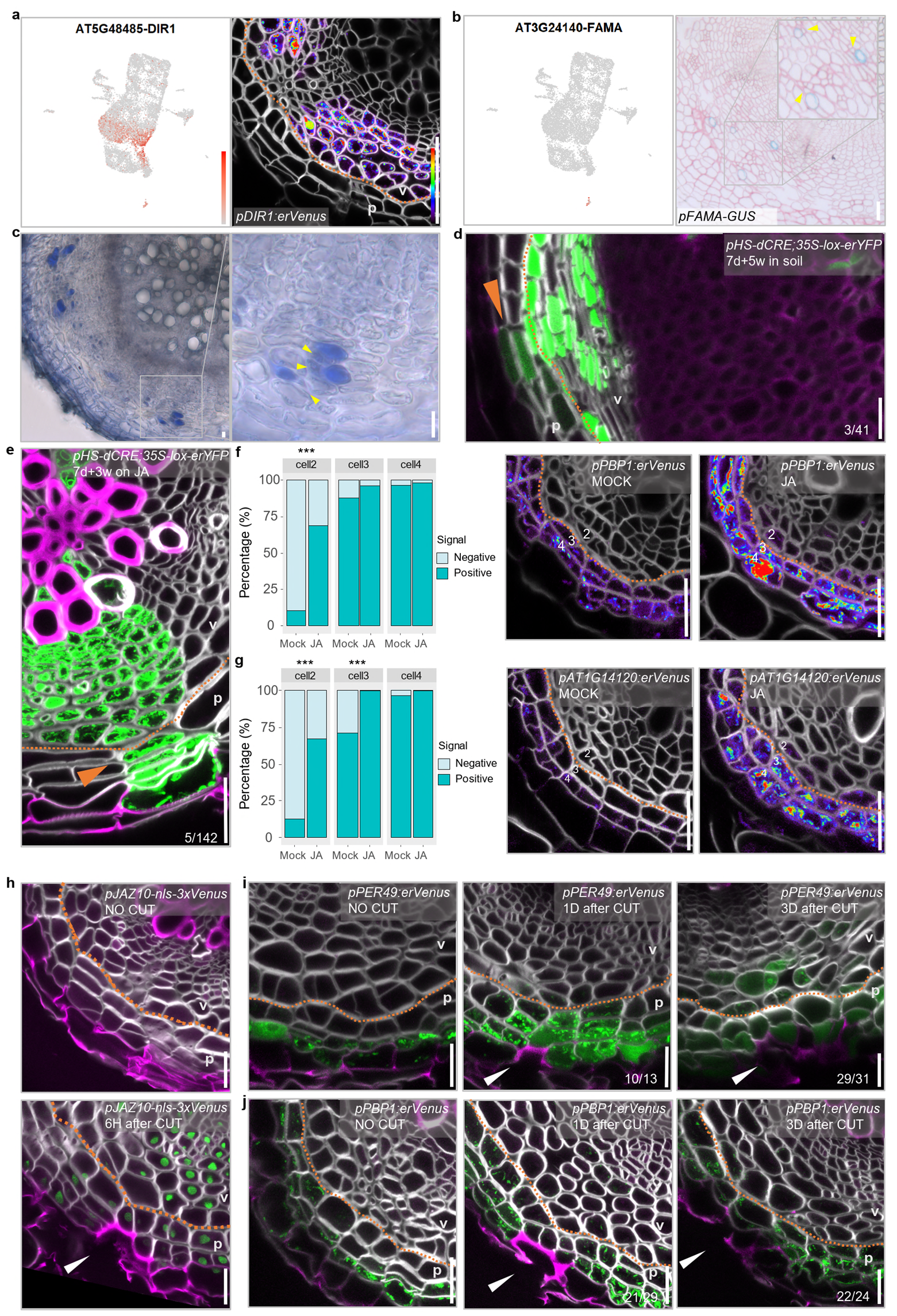
Mature phloem parenchyma functions in stress response and re-specification of the barrier upon injury. **a,** UMAP plot and confocal cross-section of 16-day-old promoter-reporter line of DIR1 which is highly detected in mature phloem parenchyma clusters, b, UMAP plot and light microscopy image of cross-section of promoter-reporter line of FAMA specifically detected in myrosin idioblast (Ml) cluster, c, Bright-field cross-section of wild-type root with CBB staining, b and c show cross­sections of 30-day-old roots. Yellow arrowheads point at sieve elements adjacent to the Mis. **d, e,** Confocal cross-section of pHS-dCRE,35S-lox-erYFP root. Clones induced in 7-day-old seedlings were analyzed 5 weeks after growth on soil (d) or 3 weeks after growth on 1/2 GM plates supplemented with 10 pM methyl jasmonate (e). Orange arrowheads indicate the transition sector. The fraction indicates 3 out of 41 (d) or 5 out of 142 (e) sectors extended to the periderm, f, g, Confocal cross-sections of 12-day-old pPBP1:erVenus **(f)** and pAT1 G14120:erVenus **(g)** roots grown without (Mock) or with (JA) methyl jasmonate for 2 days, and the frequency of signal positive cells in each cell in a cell file. Numbers in the cross-sections indicate cell position numbers in the barchart. Fisher’s exact test was performed. ***p<0.001. h, Confocal cross-sections of 17-day-old pJAZ10-nls-3xVenus roots without injury (upper) or 6 hours after the phellem and phellogen were damaged (lower), **i, j,** Confocal cross-sections of pPER49:erVenus **(i)** and pPBP1 :erVenus **(j)** roots without injury (left) and one day (middle) or three days (right) after the phellem and phellogen were damaged in 17-day-old roots. In cross sections of **a, d-j,** cell walls were stained with SR2200. In **d, e, h-j,** Green; YFP, magenta; Basic Fuchsin. In **a, d-j,** orange dashed line indicates the clonal boundary of vascular tissue and periderm., In **d, e, h-j,** v; vascular region, p; periderm. Relative expression levels of genes in UMAP and Venus signals in cross sections were shown according to the color map in **a.** In **h-j,** white arrowheads indicate the wounding site. Scale bars, 20 pm **(a-j).**

### Mature phloem parenchyma cells obtain periderm cell identity in response to superficial injury

In our dataset, the periderm clusters 12 and 3 were adjacent to but separated from mature phloem parenchyma clusters. This separation is consistent with the observation that the periderm and phloem parenchyma have a different origin in primary tissue: the periderm originates from the pericycle cells and the phloem parenchyma primarily from procambium cells^11,3^. Thus, the clonal boundary is well-defined, especially in young secondary tissues (Fig. 3a)^11^. However, we found genes which were frequently detected in both mature phloem parenchyma and periderm clusters and reporters validated expression in both tissues (Supplementary Fig. 6e). Moreover, in the dataset, there were a small group of cells between mature phloem parenchyma and periderm clusters (Fig. 1b). Since the phelloderm is adjacent to the mature phloem parenchyma, we thus hypothesized that mature phloem parenchyma cells gradually obtain the phelloderm identity. To test this hypothesis, we carried out lineage tracing by inducing YFP clones in procambial cells within a few days after the secondary growth activation. Five weeks after the induction, we examined the non-XPP lineage sectors in which the phloem parenchyma and periderm have distinct origins^11^, and found that the majority of these sectors reached the boundary of the periderm and mature phloem parenchyma (35 out of 41 sectors). However, a small proportion of sectors extended into the periderm; we found that single phelloderm cell (one cell invasion) (3 out of 41 sectors) or even an entire radial periderm cell file (one cell file invasion) (3 out of 41 sectors) was originated from the procambium (Fig. 3d). These data thus shows that mature phloem parenchyma cells have the capacity to obtain periderm identity, but at low frequency under normal growth conditions.

Next, we investigated the biological role of this transition. Mature phloem parenchyma clusters showing the gradual transition to the periderm (clusters 1 and 9), were overrepresented by jasmonic acid (JA) and salicylic acid (SA) responses and response to biotic stress (Supplementary Fig. 6f). These data suggest that mature phloem parenchyma cells might be preparing for biotic stress and these stress hormones could promote the transition of the phloem parenchyma to the periderm to reinforce the root barrier. To examine the role of JA and SA in the transition, we performed lineage tracing upon JA or SA treatment. We induced single-cell YFP clones in the procambium within a few days after the initiation of secondary growth and treated the seedlings with JA or SA for three weeks. Among 88 vascular sectors, four sectors showed one cell or one cell file invasion into the periderm upon SA treatment (one cell invasion; three out of 88 sectors, one cell file invasion; one out of 88 sectors), whereas no sector showed such transition in mock-treated roots (none out of 47 sectors) (Supplementary Fig. 7a). Upon JA treatment, five out of 142 sectors showed two cells or an entire cell file invasion into the periderm (two cell invasion; four out of 142 sectors, cell file invasion; one out of 142 sectors) (Fig. 3e). Conversely, mock-treated roots for JA treatment did not show any sectors with more than two cell invasions (75 sectors). Only three of them showed sectors with one cell invasion. These results suggest that treatment of JA and SA accelerates the transition of mature phloem parenchyma cells to periderm cells. This accelerated transition seems not to be caused by general growth retardation caused by JA and SA^34,35^ , since abscisic acid (ABA), another stress hormone and growth inhibitor^36^, did not accelerate the transition (none out of 54 sectors), despite being able to slow down radial growth. We wondered whether such a hormone treatment might affect cell identities in the mature phloem parenchyma cells as a part of the transition process. Hormonal treatment was first applied to two periderm markers, *pPBP1-erVenus* and *pAT1G14120-erVenus.* JA resulted in the most significant expansion of periderm marker expression inward, into mature phloem parenchyma. (Fig. 3f, g, Supplementary Fig. 7b). SA or ABA showed minor effect on these periderm markers (Supplementary Fig. 7b-f). We also tested JA treatment with other periderm markers generated from this study. *AT3G26450* and *PER49* reporter lines showed clear or slight expansion upon JA treatment, respectively. The reporter line of *BGLU23*, which is known to be JA induced^37^, showed drastic signal increase and expansion in the whole secondary tissues after JA treatment (Supplementary Fig. 7b,g-i). These results suggest that phloem parenchyma adjacent to the periderm obtains periderm identity as a result of JA treatment. This is in accordance with the accelerated transition as observed in our lineage tracing experiment (Fig. 3e).

Given that JA signalling is induced upon tissue injury^37^, we hypothesized that the transition is beneficial for reinforcing the barrier upon superficial injury caused by e.g. growth in soil^38^. To mimic superficial injury, we ablated the phellem and phellogen with a shallow longitudinal cut along mature roots with a razor blade. We confirmed that superficial injury was sufficient to induce JA response marker expression *pJAZ10-nls-3xVenus* near and further away from the cut site six hours after the injury (Fig. 3h). One day after the injury, the periderm markers, *PER49* and *PBP1*, expanded their expression into the mature phloem parenchyma cells, beneath the wound (Fig. 3i,j). These observations suggest an inward identity shift after the superficial injury. Re-establishment of phellogen in former phelloderm cells and the phellem cells adjacent to them were first observed three days after the injury. *PER49* and *PBP1* expression were maintained beneath the wound site to mark the newly formed periderm (Fig. 3i,j). Together, our data indicate that the phelloderm and phloem parenchyma function as a reservoir for the phellogen, and thus for the barrier re-establishment. During normal development, the transition from phloem parenchyma via phelloderm to phellogen is slow and sporadic, but this can be accelerated in case of superficial injury to reinforce the barrier.

## Discussion

In this study we used single-cell RNA sequencing and extensive reporter analysis to identify cell types and states in the *Arabidopsis* root undergoing secondary growth (Fig. 4a). By combining prior knowledge and our new findings, we were able to generate a hierarchal cell fate determination map for all cell types during secondary growth in roots (Fig. 4b; Supplementary Fig. 8). The vascular cambial stem cells produce xylem cells inwards and phloem cells outwards^11,39^. Our new data indicate that xylem-side stem cell daughters first obtain common xylem identity before specifying into one of three different cell types: vessels, parenchyma or fiber cells. Similar to the visibly differentiated xylem cell types, we found that also xylem parenchyma undergoes several maturation steps. The phloem-side stem cell daughters obtain phloem identity before becoming phloem parenchyma cells or they undergo formative divisions to form conductive phloem cells (sieve elements and companion cells) and CPP cells (Fig. 4b). We discovered that CPP cells can also be recruited from the parenchyma cell lineage by the adjacent conductive phloem cells. As vascular cambium produces more phloem cells, the previously produced phloem parenchyma cells enlarge and become mature phloem parenchyma cells. Myrosin idioblast (MI) cells are also developed within the mature phloem parenchyma region, at least in part by differentiating from mature companion cells.

**Figure 4.**
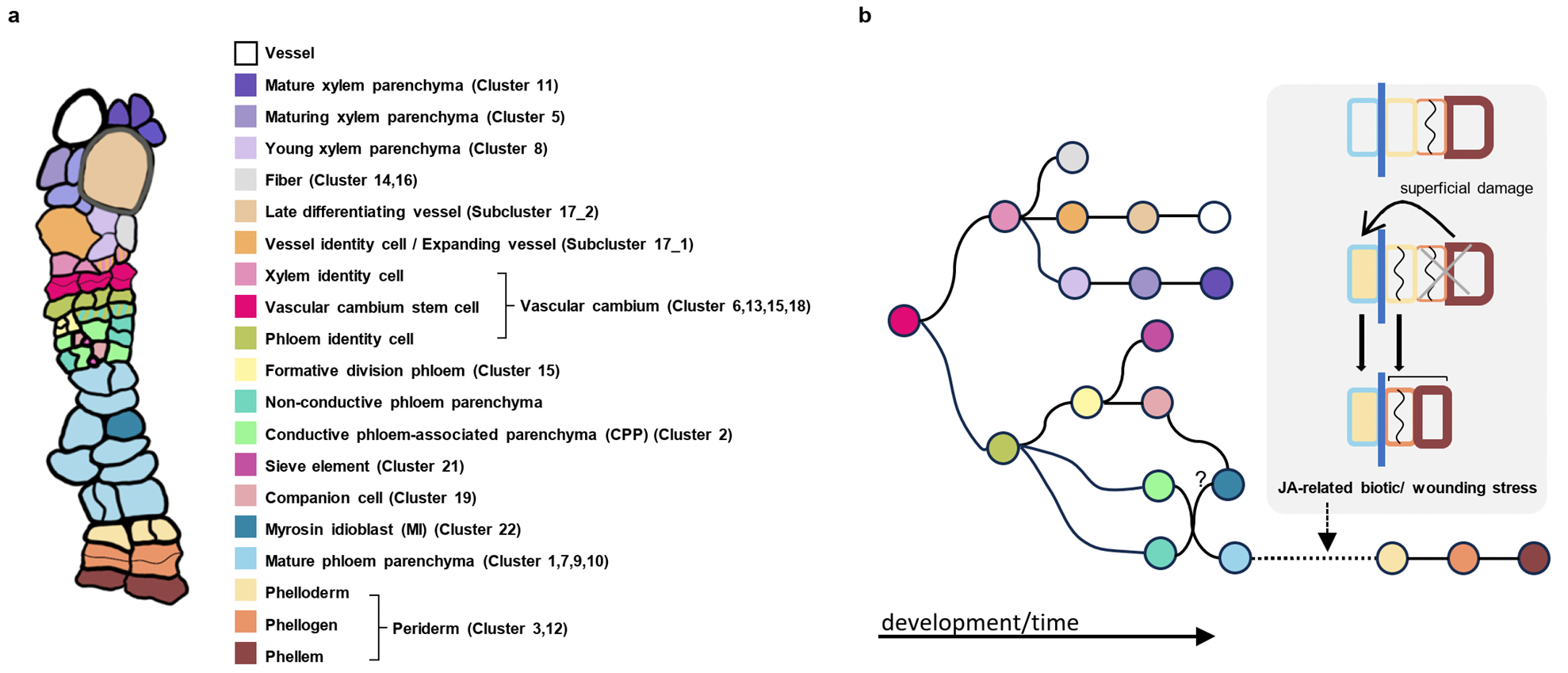
A hierarchical cell fate determination map of the *Arabidopsis* root undergoing secondary growth. **a**, Schematic illustration of cell types in Arabidopsis secondary tissues. Cell type classification based on previous reports and data presented in this study. Cluster ID corresponds to the cluster ID in the UMAP. **b**, Schematic illustration of hierarchical cell fate determination in secondary tissue. Color of each circle represents cell identity in **a**.

We made an unexpected discovery that the mature phloem parenchyma cells gradually undergo identity transition to become phelloderm, the innermost cell type of the periderm. Our data suggest that upon physical damage of the periderm, JA signalling is triggered which in turn stimulates the accelerated transition of mature phloem parenchyma into periderm cells to quickly reinforce the barrier. Since wounding caused faster cell transition than JA treatment, we hypothesize that there are also other factors than JA in promoting the transition. It has been reported that damage-induced JA signaling in root apical meristem triggers stem cell regeneration to sustain primary root growth ^40^. Whereas the underlying mechanism for the transition of phloem parenchyma into the periderm is unknown, it is possible that the common regulatory factors are utilized for stem cell regeneration in the root tip and periderm replenishment. In addition to abiotic stress, also biotic stress, such as bacteria, fungi or herbivore, can cause damage to the periderm and activate JA-induced barrier reinforcement. Additionally, herbivore attack can cause deeper damage to the secondary tissue, which can be confronted with the MI cells located within the mature phloem parenchyma. Breakage of the MI cells and the neighbouring mature phloem parenchyma cells in *Brassicaceae* such as *Arabidopsis thaliana* triggers a herbivory defence response called the mustard oil bomb^29,30^. Thus, two elaborated defence mechanisms are elicited upon injury depending on the depth of damage to protect the mature root. Protection of the mature root is critical for the plant’s survival as it connects the whole root system to the entire shoot system located above.

Storage organs consist of specialized secondary tissue produced by vascular cambium of several important crop species, such as cassava and sweet potato^41,42^. Majority of the tissue in storage organs are composed of xylem parenchyma cells, which have a function to act as storage tissue for carbohydrates and other nutrients^41,43^. Thus, our parenchyma transcriptome dataset will be valuable in identifying state-specific roles of parenchyma cells, which may have a function in corresponding storage cell types in crop species.

## Methods

### Plant growth condition

The seeds were sterilized with 20% chlorine for 3 min followed by sterilization with 70% ethanol for 5 min and were washed with milliQ water twice. The seeds were kept in dark at 4 °C for two days and then plated on half-strength Murashige and Skoog growth medium (1/2 GM) supplemented with 2.2 g/L MS salt mixture with vitamins (Duchefa), 1% sucrose, 0.5 g/L MES and 0.8% agar (pH 5.8). In case of the T1 generation seeds, the seeds were mixed with 0.1% agar supplemented with 250 µg/ml Cefotaxime (Duchefa) before plating to avoid Agrobacterium contamination. The plates were placed vertically in a growth chamber. The day when the plates were moved to the growth chamber was defined as day 0. Analysis was performed with plate-grown seedlings unless otherwise mentioned. The seedlings were transferred to soil around day 7 if necessary for analysis. The plants were grown at 23 °C under long-day photoperiod conditions (16 hours light/8 hours dark).

### Protoplast isolation and florescence-activated cell sorting

We utilized 30-day-old *pPXY:erYFP* (Col-0) grown on soil to isolate protoplasts for single cell analysis. The protoplast isolation protocol was modified from the published protocol ^44^. Roots within 2 cm below the root-hypocotyl junction were harvested (lateral roots were removed) and washed with tap water and then Milli Q water. The roots were longitudinally dissected with a razor blade under a stereo microscope and put into the protoplast isolation solution (1.5% (w/v) cellulase-R10 (Yakult), 0.4% (w/v) macerozyme-R10 (Yakult), 0.4 M mannitol (Sigma-Aldrich), 20 mM MES (Duchefa), 20 mM KCl (1 M stock in Milli-Q water), 0.1% (w/v) BSA (Sigma-Aldrich) and 10 mM CaCl2 (1 M stock in Milli-Q water)). The samples were incubated at room temperature under the dark condition with gentle shaking (75 rpm) for one hour. After the incubation, the solution was filtered once with the 70 µm cell strainer and the flow-through was centrifuged at 400 g for 6 min. The supernatant was gently removed, and the protoplasts were resuspended in the buffer (the protoplast isolation solution without enzymes). The resuspended protoplast solution was filtered three times with the 40 µm cell strainer. Protoplasts were stained with 14 µM 4’,6-diamidino-2-phenylindole (DAPI) in PBS and the DAPI negative cells were sorted by BD FACSAria II. We confirmed the efficiency of this protocol by isolating protoplasts from known tissue-specific reporters covering the major cell types in the root secondary tissue. We were able to capture fluorescence-positive protoplasts from each reporter lines, suggesting that our protoplast isolation method was sufficient to acquire cells residing in different regions of the secondary tissue.

### Single-cell RNA sequencing sample processing, library establishment and sequencing

After sorting, the protoplasts were centrifuged at 400 g at 4°C for 5 minutes and then resuspended in resuspension solution to a final concentration of around 1,000 cells/µL. Resuspended cells were loaded on a Chromium Single Cell 3’ GEM, Library & Gel Bead Kit (V3 chemistry, 10X Genomics) according to the manufacturer’s instructions. Libraries were sequenced on an Illumina HiSeq4000 and NovaSeq6000 instrument following recommendations of 10X Genomics at the VIB Nucleomics Core facility (VIB, Leuven).

### Processing of raw sequencing data and data analysis

The FASTQ files obtained after demultiplexing were used as the input for ‘cellranger count’ (version 6.1.2), with reads being mapped to the *Arabidopsis thaliana* reference genome (Ensembl TAIR10.40). Initial filtering in CellRanger recovered 17,140 cells, corresponding with 33,679 mean reads per cell and 2,409 median genes per cell. Further data processing was performed in R (version > 3.6.0) (https://www.r-project.org/) by the ‘scater’ package^45^ (version 1.10.1). Outlier cells were defined as having less than 4,000 UMIs or as cells containing more than 5% mitochondrial or chloroplast transcripts. After removing outliers, 11,760 cells were retained for further analysis. Normalizing the raw counts, detecting highly variable genes, finding clusters and creating Uniform Manifold Approximation and Projection (UMAP) plots were done using the Seurat package (version 4.1.0)^46^. Differential expression analysis for marker gene identification per subpopulation was based on the non-parametric Wilcoxon rank sum test implemented within the Seurat pipeline. Necessary reported information to allow evaluation and repetition of a plant single cell/nucleus experiment is included in Supplementary Table S1.

### Gene ontology (GO) enrichment analysis

Differentially expressed gene (DEG) list (avg_logF2C ≥ 0.5) of each cluster or subcluster was used for Gene ontology (GO) enrichment analysis using the clusterProfiler v 4.2.2^47^ in the program R v4.0.2 (https://www.r-project.org/) with p < 0.05.

### Cloning of reporter lines and plant transformation

We selected genes for cluster validation based on their expression in clusters in the dataset. Their promoter regions were amplified with primers listed in Supplementary Table S4 and cloned into the *pDONRP41R* entry vector as the first box by Gateway BP Clonase II enzyme (Thermo Fisher Scientific). The first box plasmid (promoter), the second box plasmid (reporter gene), *221z-erVen* or *221z-erRFP*^48^, the third box plasmid (terminator), *2R3e-3AT*^48^, and the destination vector, *pFRm43GW*^49^ containing an *RFP* seed coat selection marker, were used to generate the reporter line constructs by MultiSite Gateway technology (Thermo Fisher Scientific). The genes selected for the reporter analysis, the primers used for promoter amplification, and the frequency of T1 individuals showing expression pattern predicted from scRNA-seq dataset and the available generation of the reporter seeds are listed in Supplementary Table S4. The generated promoter:*erVenus-YFP* constructs were transformed into Col-0 and the reporter expression was examined in T1 generation unless otherwise stated. The *pTET5:erRFP* was transformed in *HS-dCRE;35S-lox-erYFP*.

### Mutant analysis

In this study, Col-0 was used as wildtype. The *3paplC* triple mutant was received from authors reported the mutant^19^. The rest of the mutants were ordered from Nottingham Arabidopsis Stock Centre (NASC): *apk1a* (GK-430G06), *at4g30460* (SALK_140721), *anac087* (SALK_079821), *myb3r-3* (SALK_143357C), *myb3r-5* (SALK_205058C), *nac055* (SALK_014331C), *myb112* (SALK_017020C), *anac046* (SALK_107861C), *at4g21620* (SALK_099390C). The mutants ordered from NASC were genotyped and homozygous lines were selected for seed propagation. All the mutants and Col-0 were freshly propagated for phenotyping, The mutant phenotypes were examined utilizing 14-day-old seedlings except for *3paplC* which were 16-day-old. The phenotyping of *3paplC* were repeated four times. The phenotype of the other mutants was repeated twice.

### Lineage tracing analysis

For lineage tracing analysis, we generated transgenic plants in which *er-YFP* sectors can be activated upon heat-shock based on previously generated line^11^. We backcrossed *HS-dCRE;35S-lox-GUS* line with Col-0 and established the F3 line which was homozygous for *HS-dCRE* without *35S-lox-GUS* construct (*HS-dCRE*) based on selection markers (hygromycin for *HS-dCRE* and Basta for *35S-lox-GUS*). The *35S-lox-erYFP* construct was transformed into the *HS-dCRE* seedlings; the T3 line homozygous for *35S-lox-erYFP* construct (*HS-dCRE;35S-lox-erYFP*) were established based on the selection marker (Basta). T3 or T4 seedlings were utilized for analysis. *pTET5:erRFP* was transformed into *HS-dCRE;35S-lox-erYFP* and were analysed at T2 generation.

For clone induction, 7-day-old seedlings grown on 1/2GM plates were utilized. After removing excess water from plates, one or two plates were placed in a plastic bag. We sealed the plastic bag with plastic tape to avoid water leakage and submerged the bag in 37°C water for 20 min. We retrieved plates from the plastic bag and horizontally kept the plates at 4°C for 30 min. The plates were moved to the growth chamber. One day later, the sector induction was examined under a fluorescence stereo microscope. For sector analysis at five weeks after the induction, the seedlings were moved to soil. For sector analysis with hormone treatment, seedlings were transferred to 1/2GM plates supplemented with hormone.

### Chemical treatment

For hormone treatment, salicylic acid (SA, Sigma-Aldrich), methyl jasmonate (JA, Sigma-Aldrich), and abscisic acid (ABA, Duchefa) were dissolved in DMSO (SA and JA) or ethanol (ABA) to prepare 100 mM stock solution. The stock solution was stored at −20 °C. To examine gene expression upon treatment, the homozygous T3 reporter lines were first grown on 1/2 GM plates for 10 days, then were transferred to 1/2 GM plates supplemented with 5 µM SA, 10 µM JA or 10 µM ABA, or 1/2 GM plates contained an equal volume of DMSO or ethanol as a control (called Mock in the experiments). The seedlings were treated for 2 days.

For lineage tracing analysis with hormone treatment, 7-day-old seedlings grown on 1/2 GM plates were utilized to induce clones upon heat as described above. One day later, the seedlings were transferred to 1/2 GM plates supplemented with 5 µM SA, 10 µM JA or 10 µM ABA, or 1/2 GM plates contained an equal volume of DMSO or ethanol as a control (Mock). The seedlings were grown for 20 days.

The 17-β-oestradiol (EST, a synthetic derivative of oestrogen, Sigma-Aldrich) was prepared as 20 mM stock in DMSO and the stock solution was stored at -20°C. The 13-day-old *p35S: XVE>>CKX7*^50^ seedlings were transferred to the 1/2 GM plate containing 5 µM EST or equal amount of DMSO(Mock) for 1 day.

### Fluorescence marker fixation, vibratome sectioning and confocal imaging

The protocol was modified from published paper^11^. To examine fluorescence reporter expression, samples were fixed in 4% paraformaldehyde (PFA, Sigma-Aldrich) in 1×phosphate-buffered saline (PBS, pH 7.2) supplemented with 0.1% triton under vacuum for 1h, then washed twice with 1xPBS. Roots were embedded into 5% agarose in 1xPBS, followed by vibratome sectioning. The 200 µm thick slices were kept in 1xPBS supplemented with 1 µl/ml SR2200 (Renaissance Chemicals) to stain cell wall. To stain lignin, the agarose slices were put into ClearSee^51^ supplemented with 1 µl/ml SR2200 and 50 µg/ml Basic Fuchsin (Sigma-Aldrich)^52^. The cross sections were mounted with 1xPBS or ClearSee and imaged with Leica SP8 Stellaris confocal microscope (Leica) under 20x or 63× objectives with Leica Las AF software. All the confocal images were taken with sequential scan mode. For *p35S:XVE>>CKX7*^50^ , the sections were stained with 1 μg/ml calcofluor white (Sigma-Aldrich) in 1xPBS. To visualize reporter gene expression, the settings were adjusted individually for each reporter line. However, when fluorescent signal intensities were compared within an experiment, identical settings were used. For the setting to visualize cell wall staining, signals were adjusted for each cross section individually.

### GUS staining, material fixation, microtome sectioning and light microscopy

The GUS staining protocol was modified from published paper^53^. Samples were kept in 90% acetone on ice for 30 min. After the incubation, the samples were washed twice with 0.05 M sodium phosphate buffer (pH 7.2). Then the samples were submerged into GUS solution (0.05 M sodium phosphate buffer (pH 7.2), 1.5 mM ferrocyanide, 1.5 mM ferricyanide, 1 mM X-glucuronic acid and 0.1% Triton X-100) and kept at room temperature under vacuum for 1h. The samples were incubated at 37 °C until the desired GUS signals were detected.

Samples for microtome sectioning were fixed in 1% glutaraldehyde and 4% formaldehyde in 0.05 M sodium phosphate (pH 7.2) for overnight and washed with 0.05 M sodium phosphate twice. To gradually dehydrate, the samples were kept in 10%, 30%, 50%, 70%, 96% and 100% ethanol for 30 min in each step. The 30 min incubation with 100% ethanol was repeated one more time. The samples were transferred to a 1:1 (v/v) solution of 100% ethanol and solution A (Leica Historesin Embedding kit) and kept for 1h, followed by the incubation in solution A overnight. The samples were aligned in chambers and embedded with a 14:1 (v/v) solution of solution A and the hardener.

The cross sections of embedded samples were made with Leica JUNG RM2055 microtome. The sections were acquired around 0.5 cm below the root-hypocotyl junction. The thickness of the sections was 10 µm for GUS reporter lines or otherwise 5 µm. The cross sections of GUS reporter lines were staining with 0.05% (w/v) ruthenium red (Sigma-Aldrich); the cross sections without GUS signals were additionally stained with 0.05% (w/v) toluidine blue (Sigma-Aldrich). The sections were imaged with Leica 2500 microscope under 20× and 40× objectives.

### CBB staining

The protocol was modified from published paper^33^. Thirty-day-old plants were collected in 15 mL CELLSTAR tube (Greiner Bio-One) with CBB solution (45% methanol, 10% acetic acid, and 0.25% CBB R250) and boiled in water bath for 3 min, followed by wash with 1xPBS three times. The roots within 1 cm below the root-hypocotyl junction were embedded in 5% agarose and sectioned by vibratome with 100 µm thickness. The sections were mounted with water and imaged with Leica 2500 microscope.

### Wounding experiment

We used 17-day-old seedlings for the cutting experiment. Roots within 5 mm from the root-hypocotyl junction were longitudinally injured with a razor blade under a dissection stereo microscope. We defined the superficial cut as the ablation of the phellem and phellogen. Only cross sections with the superficial cut were used for analysis.

### Quantification and statistical analysis

For mutant phenotype quantification, number of secondary vessels, and sieve element, vasculature diameter and cellular fluorescence signal intensity were manually measured by FIJI ImageJ v1.52^54^. All the plots for visualizing the quantification results were produced by ggplot2 package^55^ in RStudio (https://www.rstudio.com/) with the program R v4.0.2 (https://www.r-project.org/). For boxplots, the boxes in the box-and-whisker plots represent median values and interquartile range, the whiskers indicate the total range. The black dots indicate measurements from individual roots. For each reporter line, fluorescence signal intensities were measured in cell1 to cell4 (Supplementary Fig. 5b). We defined signal ‘positive’ cells as cells in cell2, 3, 4 whose signal intensities were higher than maximum signal intensity in cell1 of mock-treated roots. The proportion of signal positive cells in cell2, 3, and 4 was visualized by bar charts.

For mutant phenotype analysis, the normality distributions of each type of quantification were tested with Shapiro-Wilk normality test. Significant differences were examined by two-tailed Wilcoxon test, except for number of sieve element, of which significant test was performed by two-tailed *t*-test. For fluorescence signal analysis, the Fisher’s exact test was performed.

## DATA AND MATERIALS AVAILABILITY

The data can be accessed via a freely accessible on-line browser tool (https://www.single-cell.be/plant) The transcriptome data is deposited in the online tool and raw data can be accessed at NCBI with GEO number: GSE270140. All other data are either in the main paper or the Supplement. Material requests should be directed to the corresponding authors.

## ACKNOWLEDGEMENT

We thank Jos Wendrich for helping the sample processing of single-cell RNA sequencing and Kevin Verstaen for helping data processing in VIB; Ykä Helariutta, Sofia Otero, Pawel Roszak for providing published seeds; Xixi Zhang, Jennifer López Ortiz, Tiina Blomster, Tonni Grube Anderson and Ryohei Thomas Nakano providing suggestions to the manuscript; Dominique C Bergmann for providing the pFAMA-GUS made by Anne Vatén during her post-doctoral period; Kenneth Birnbaum for providing suggestion for protoplast isolation and data analysis; special thanks to Mikko Herpola and Miki Iida for laboratory management and Light Microscopy Unit (LMU), University of Helsinki, for providing the confocal microscopy equipment and technical assistance.

This work was supported by the Research Council of Finland (grant numbers 316544 and 346141 to M.L., H.I., M.M., S.M., L.Y., B.W., X.W. and A.P.M.), European Research Council (ERC-CoG CORKtheCAMBIA agreement 819422 to M.L., H.I., L.Y., B.W., X.W. and A.P.M.), University of Helsinki (Doctoral Programme in Plant Biology to M.L.), an EMBO Postdoctoral Fellowship (ALTF 128-2020 to H.I.), the Japan Society for the Promotion of Science (Overseas Research Fellowships to H.I.), and the European Research Council (ERC StG TORPEDO; 714055 and ERC CoG PIPELINES; 101043257 to T.E. and B.D.R.).

## CONTRIBUTIONS

A.P.M. conceived the project; A.P.M., M.L. and H.I. designed the experiments; B.D.R. supervised the single cell RNA-seq data analysis; M.L., H.I. and M.M. performed the experiments; T.E. performed the sequencing data analysis; S.M. integrated the dataset to Cella model; L.Y., A.V. and X.W. helped with cloning; B.W. helped with reporter analysis. A.P.M., M.L., H.I. and B.D.R. wrote the paper with input from all authors.

## COMPETING INTERESTS STATEMENT

Authors declare that they have no competing interests.

---

**Supplementary Figure 1.**
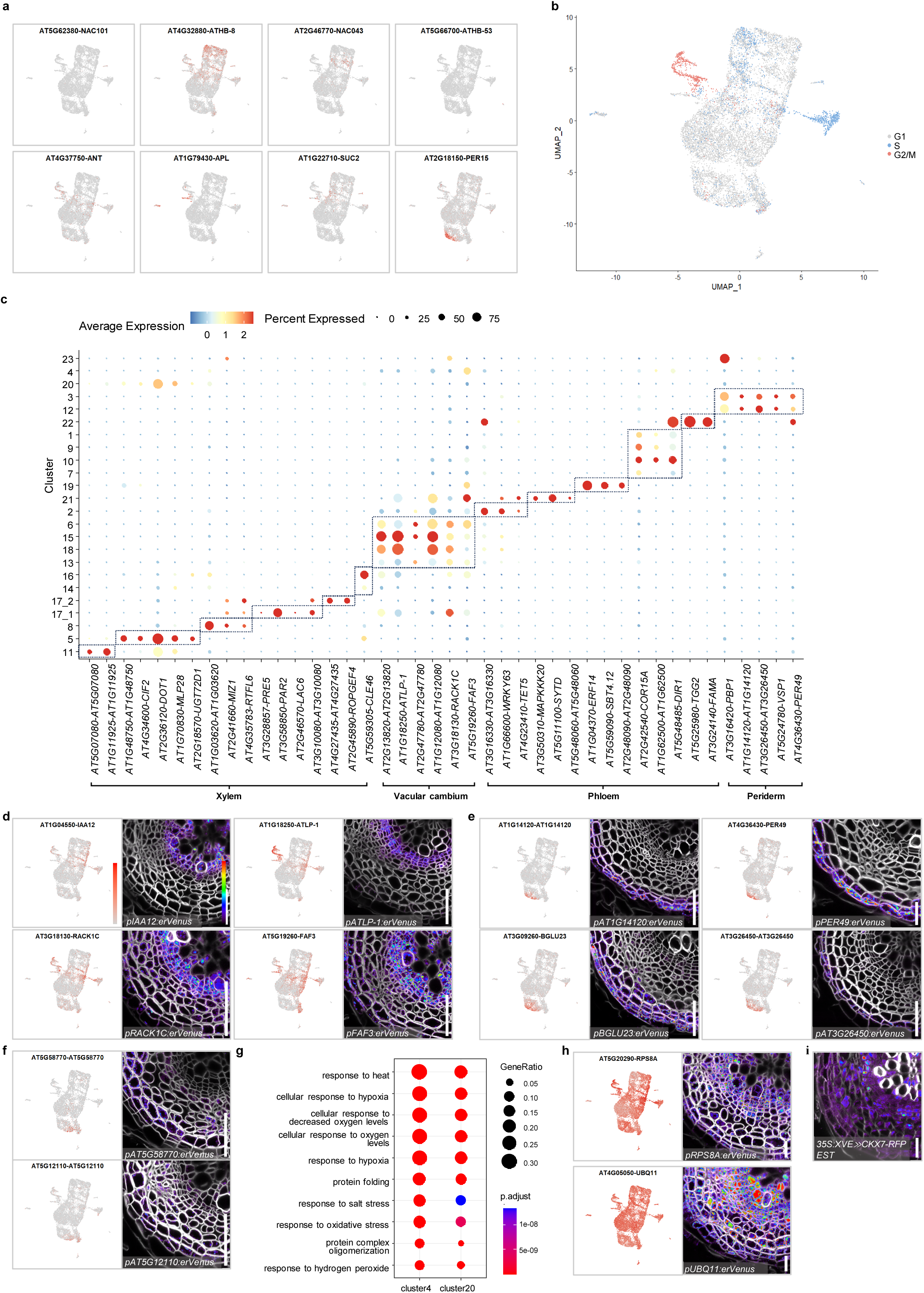
An overall analysis of the single cell RNA-seq data. **a**, UMAP plots of known tissue-specifically expressed genes (*NAC101, ATHB8, NACO43, ATHB53, ANT, APL, SUC2*, and *PER15*). **b**, Predicted classification of each cell in either G2/M, S or G1 phase in UMAP. Cell cycle list from Zhang et al, 2019 was utilized. **c**, Dotplot of expression of representative marker genes used for cluster annotation and data validation. Size or colour of dots represents the percentage of cells expressing a given gene or the scaled average expression level, respectively. Box with dotted line represents tissue types. **d**, UMAP plots of genes (*IAA12, ATLP-1, RACK1C* and *FAF3*) highly detected in vascular cambium clusters and confocal cross-sections of their transcriptional reporter lines. **e**, UMAP plots of genes (*AT1G14120, PER49, BGLU23* and *AT3G26450*) highly detected in periderm clusters and confocal cross-sections of their transcriptional reporter lines. **f**, UMAP plots of genes (*AT5G58770* and *AT5G12110*) highly detected in cluster 4 and 20, and confocal cross-sections of their transcriptional reporter lines. **g**, Gene Ontology (GO) enrichment analysis and comparison of differentially expressed gene (DEG)s of cluster 4 and 20. Top 10 enriched biological processes are shown. Dot size represents percentage of DEGs in a given cluster involved in this biological process; dot color indicates P.adjust value of each biological process in a given cluster. The full list of GO terms of all the clusters are available in Supplementary Table S3. **h**, UMAP plots of genes (*RPS8A* and *UBQ11*) broadly and highly detected in all clusters and confocal cross-sections of their transcriptional reporter lines. **i**, Confocal microscopy of cross-section of 13-day-old *35S:XVE>>CKX7-RFP* root grown with 5μM estrodiol (EST) for 1day. Cell wall stained with calcofluor was shown in grey. In **d**-**f**, for each gene, the UMAP plot was shown in left and cross section was shown in right. **d**-**f** and **h** show confocal cross-sections of 16-day-old roots. Cell walls stained with SR2200 were shown in grey. Relative expression levels of genes in UMAP and Venus-YFP or RFP signals in cross sections were shown according to the color map in **d**. Scale bars, 20 µm (**d**-**f**, **h,i**).

**Supplementary Figure 2.**
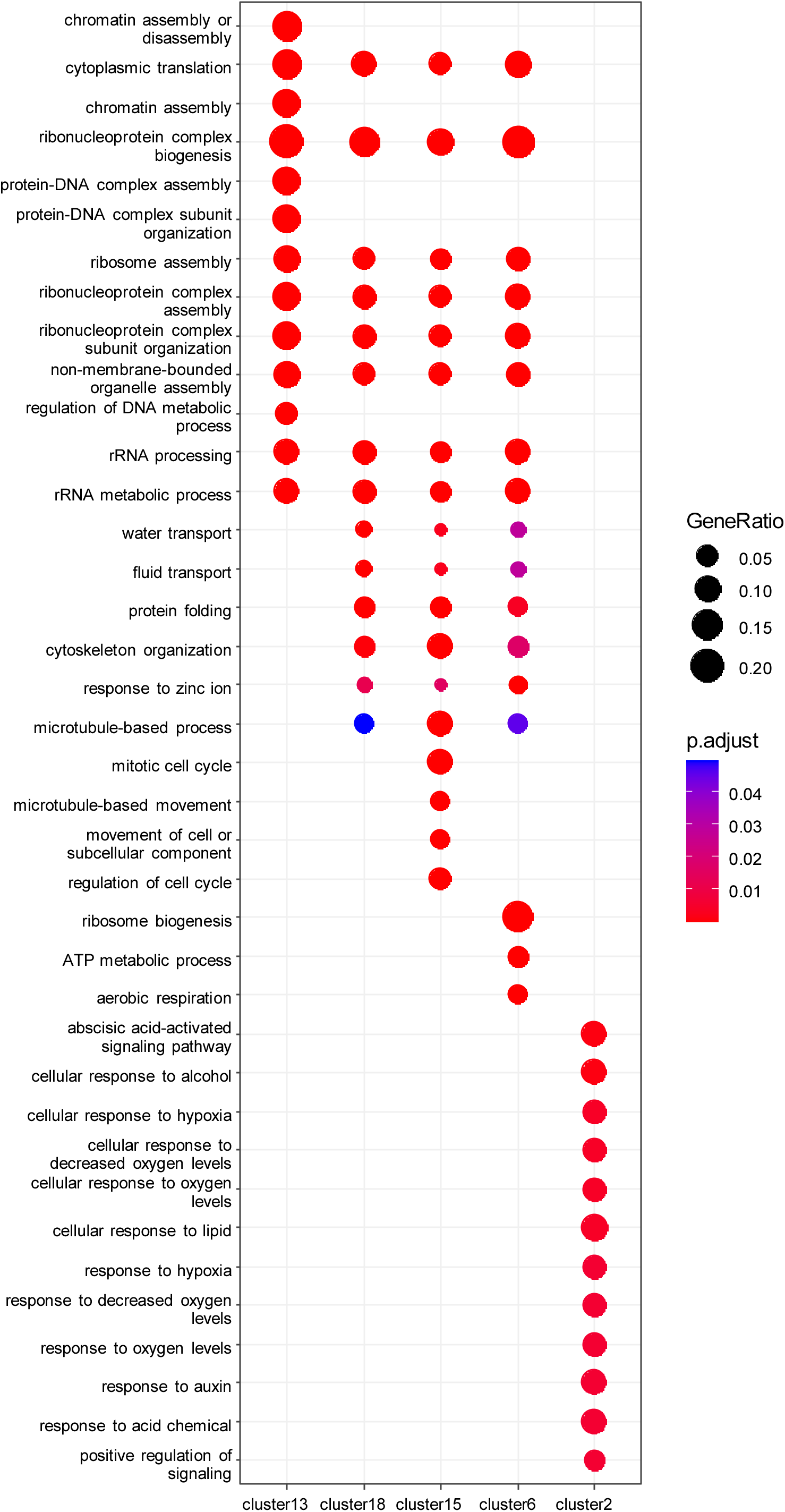
GO term comparison among vascular cambium clusters. GO enrichment analysis and comparison among DEGs of cluster 2, 6, 13, 15 and 18. Top 12 enriched biological processes are shown. Dot size represents percentage of DEGs in a given cluster involved in the biological process; dot color indicates P.adjust value of each biological process in a given cluster. The full list of GO terms of all the clusters are available in Supplementary Table S3.

**Supplementary Figure 3.**
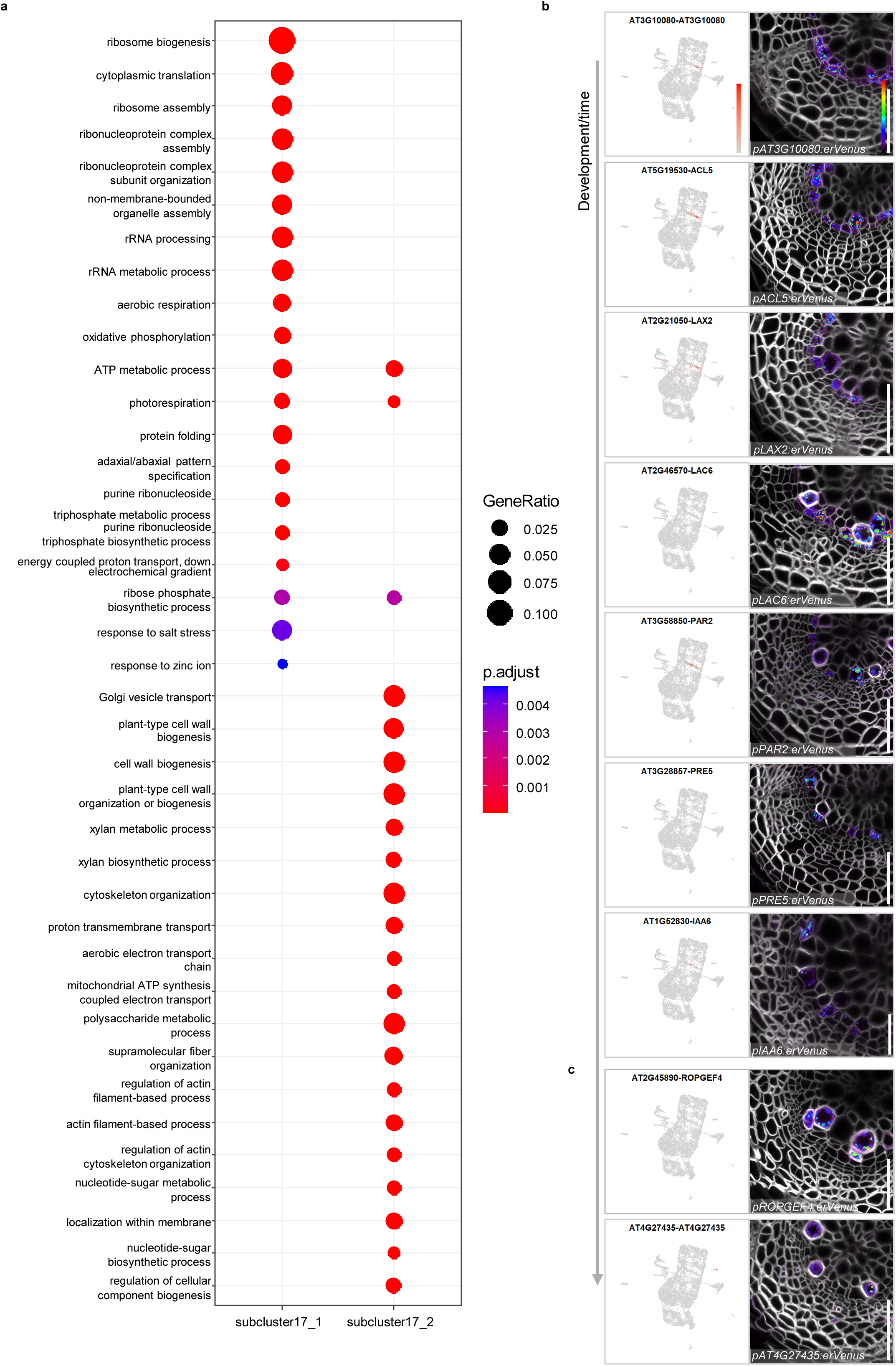
Distinct states during vessel development. **a**, GO enrichment analysis and comparison of DEGs of subcluster 17_1 and 17_2. Dotplot shows top 20 enriched biological processes. Dot size represents percentage of DEGs in a given cluster involved in the biological process, dot color indicates P.adjust value of each biological process in a given cluster. The full list of GO terms of both subclusters are available in Supplementary Table S3. **b**, UMAP plots of genes (*AT3G10080*, *ACL5, LAX2, LAC6, PAR2, PRE5* and *IAA6*) specifically detected in vessel identity cell / expanding vessel cluster and confocal cross-sections of their transcriptional reporter lines. **c**, UMAP plots of genes (*ROPGEF4* and *AT4G27435*) specifically detected in late differentiating vessel cluster and confocal cross-section of their transcriptional reporter lines. In **b**, **c**, for each gene, the UMAP plot was shown in left and cross section of 16-day-old root was shown in right. Cell walls stained with SR2200 were shown in grey. Relative expression levels of genes in UMAP and Venus signals in cross sections were shown according to the colour map in **b**. Scale bars, 20 µm (**b**, **c**).

**Supplementary Figure 4.**
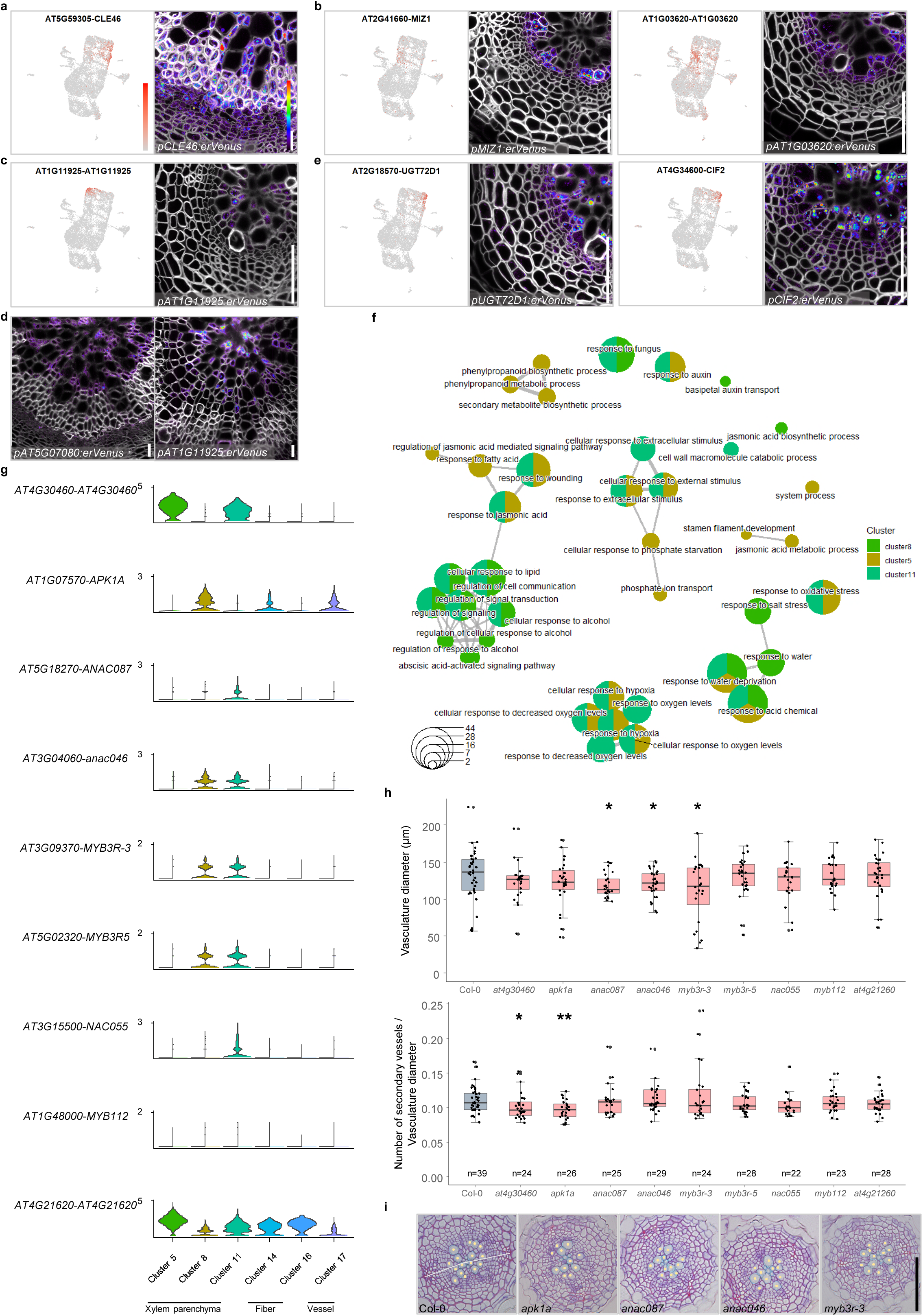
Analysis of xylem fiber and parenchyma clusters. **a**, UMAP plot of *CLE46* highly detected in differentiating fiber clusters and confocal cross-section of its transcriptional reporter line. **b**, UMAP plots of genes (*MIZ1* and *AT1G03620*) highly detected in young xylem parenchyma cells and confocal cross-sections of their transcriptional reporter lines. **c**, UMAP plot of *AT1G11925* highly detected in mature xylem parenchyma cells and confocal cross-sections of its transcriptional reporter line. **d**, Confocal cross-sections of transcriptional reporter lines (*AT5G07080* and *AT1G11925*). **e**, UMAP plots of genes (*UGT72D1* and *CIF2*) highly detected in maturing xylem parenchyma cells and confocal cross-sections of their transcriptional reporter lines. In **a**-**c** and **e**, for each gene, the UMAP plot was shown in left and cross section of 16-day-old (**b**, **c**, **e**) and 30-day-old (**a**, **d**) roots was shown in right. In cross sections of **a**-**e**, cell walls stained with SR2200 were shown in grey. Relative expression levels of genes in UMAP and Venus signals in cross sections were shown according to the colour map in **a**. **f**, GO enrichment analysis and comparison among DEGs of cluster 5, 8 and 11 shown with emapplot. The top 15 enriched biological processes were shown. Dot size represents the number of DEGs in a given cluster involved in the biological process, dot color of the cluster correlates to the cluster color in the UMAP. The edge represents the similarity between terms, shorter and thicker edge indicates higher similarity. The full list of GO terms of all clusters is available in Supplementary Table S3. **g**, Violin plots showing of genes highly expressed in xylem parenchyma clusters. The height and width of the plots represent the expression level **and** the proportion of cells showing expression, respectively. **h**, Quantification of vasculature diame ter and ratio between secondary vessels and vasculature diameter in 14-day-old seedlings in which each gene shown in **g** was mutated. The boxes in the box-and-whisker plots represent median values and interquartile range, the whiskers indicate the total range. The black dots indicate measurements from individual roots. Shapiro-W ilk normality test followed by a two-tailed Wilcoxon test was utilized for statistic test between Col-0 and mutants. * *p*<0.05, \*\**p*<0.01. The experiment was repeated twice; n indicated number of individuals analysed. **i**, Bright-field cross sections of 14-day-old wildtype (Col-0), *apk1a*, *anac087, anac046*, and *myb3r*-3 single mutant roots. Yellow dots indicate secondary vessels. White line indicates the vasculature diameter based on primary xylem and two primary phloem poles. Scale bars, 20 µm (**a**-**e**), 50 µm (**i**).

**Supplementary Figure 5.**
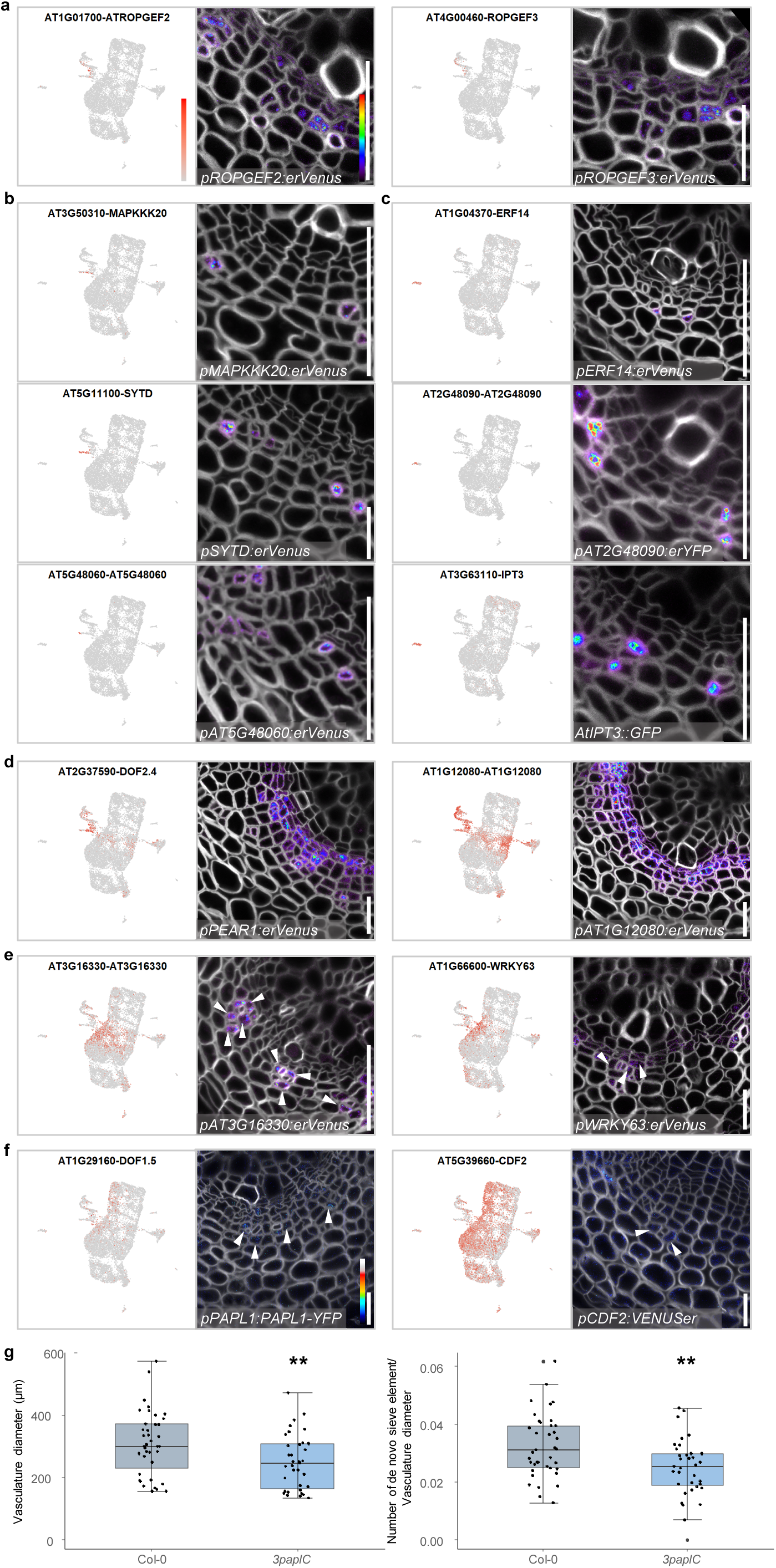
Analysis of phloem clusters. **a**, UMAP plots of genes *(ROPGEF2* and *ROPGEF3*) highly detected in formative division phloem clusters and confocal cross-sections of their transcriptional reporter lines. **b**, UMAP plots of genes (MAPKKK20, *SYTD* and *AT5G48060*) specifically detected in differentiating sieve element cluster and confocal cross-sections of their transcriptional reporter lines. **c**, UMAP plots of genes (*ERF14*, *AT2G48090* and *IPT3*) specifically detected companion cell cluster and confocal cross-sections of their transcriptional reporter lines. **d**, UMAP plots of genes (*DOF2.4*/*PEAR1* and *AT1G12080*) highly detected in phloem-side cambium clusters and confocal cross-section of their transcriptional reporter lines. **e**, UMAP plots of genes (*AT3G16330* and *WRKY63*) highly detected in CPP cluster and confocal cross-sections of their transcriptional reporter lines. **f**, UMAP plots of *PAPL* genes (*DOF1.5*/ *PAPL1* and *CDF2*) and confocal cross-sections of their reporter lines. Note: expression of CDF2 is broader than just cluster 2 in UMAP. Relative expression levels of Venus signals in cross sections were shown according to the colour map. In **e**, **f**, white arrowheads indicate CPP cells. In **a-f**, for each gene, the UMAP plot was shown in left and cross section was shown in right. **a-f** show confocal cross-sections of 16-day-old roots. Cell walls stained with SR2200 were shown in grey. Relative expression levels of genes in UMAP (**a**-**f**) and fluorescence signals in cross sections (**a**-**e**) were shown according to the colour map in **a**. **g**, Quantification of vasculature diameter and ratio between number of *de novo* sieve elements and vasculature diameter of 16-day-old wildtype and *3paplC* roots. The boxes in the box-and-whisker plots represent median values and interquartile range, the whiskers indicate the total range. The black dots indicate measurements from individual roots. Shapiro-Wilk normality test followed by a two-tailed Wilcoxon test was utilized for statistic test between Col-0 and *3paplC*. \**p*<0.05, \*\**p*<0.01. The experiment was repeated four times, thirty-nine wildtype seedlings and thirty-five *3paplC* seedlings were examined in total. Scale bars, 20 µm (**a-f**).

**Supplementary Figure 6.**
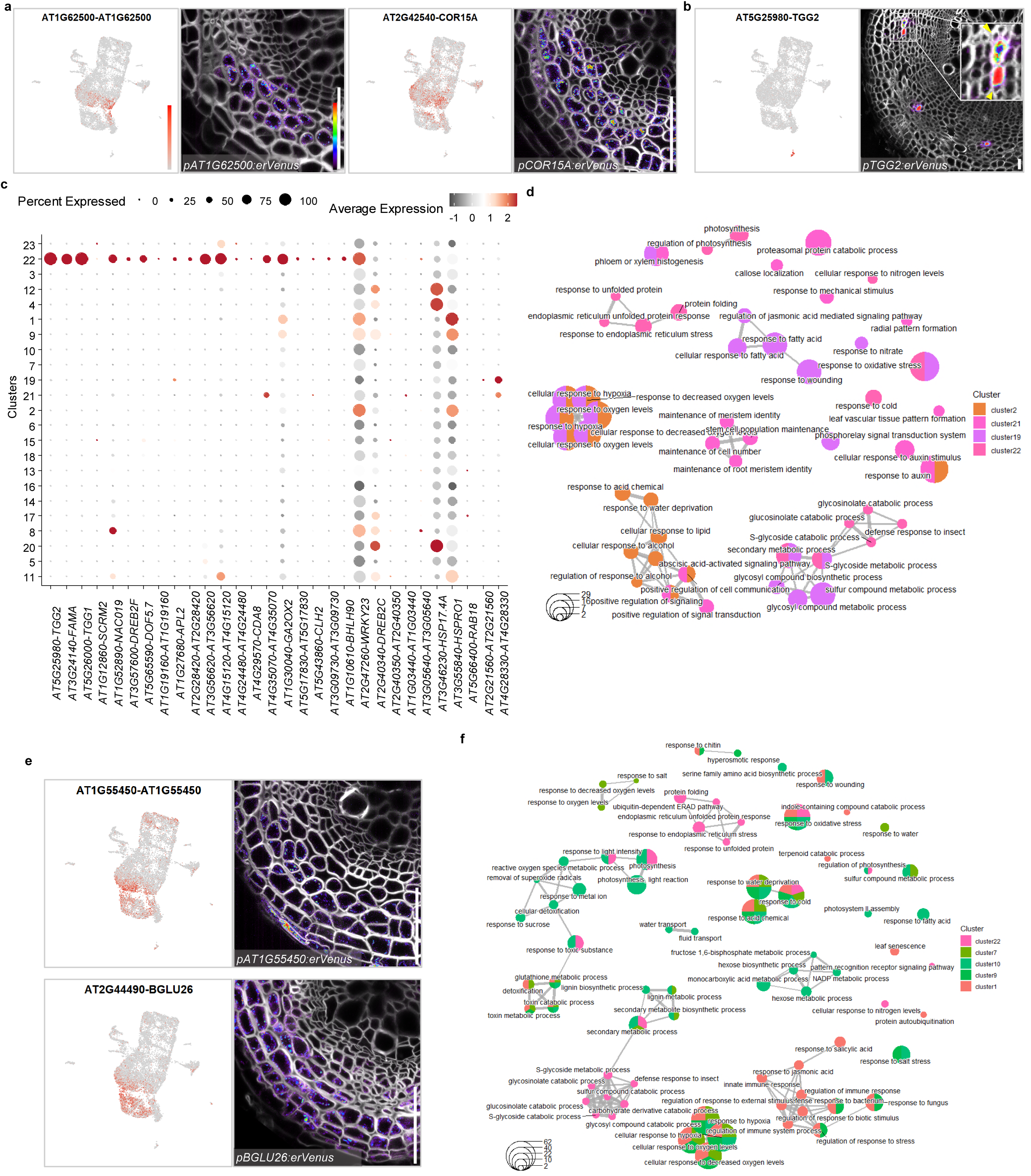
Analysis of mature phloem parenchyma clusters. **a**, UMAP plots of genes (*AT1G62500* and *COR15A*) highly detected in mature phloem parenchyma clusters and confocal cross-sections of their transcriptional reporter lines. **b**, UMAP plot of *TGG2* specifically detected in MI cluster and confocal cross-section of its 30-day-old reporter line root. Yellow arrows point at the sieve elements adjacent to the MIs. **c**, Expression of potential MI differentiation regulators in our scRNA-seq data. Cluster 22 consists of the MI cells. Dot size represents the percentage of cells expressing a given gene; colour of the dot indicates the scaled average expression level. **d**, GO enrichment analysis and comparison among DEGs of cluster 2, 19, 21 and 22 shown with emapplot. The top 15 enriched biological processes were shown. **e**, UMAP plots of genes (*AT1G55450* and *BGLU26*) highly detected in part of mature phloem parenchyma clusters and periderm clusters and confocal cross-sections of their transcriptional reporter lines. **f**, GO enrichment analysis and comparison among differentially DEGs of cluster 1, 7, 9, 10 and 22 shown with emapplot. The Top 30 enriched biological processes were shown. **a** and **e** show confocal cross-sections of 16-day-old roots. In cross sections of **a**, **b**, **e**, cell walls stained with SR2200 were shown in grey. For each gene, the UMAP plot was shown in left and cross section was shown in right. Relative expression levels of genes in UMAP and Venus signals in cross sections were shown according to the color map in **a**. In **d** and **f**, dot size represents number of DEGs in given cluster involved in this biological process; dot color of the cluster correlates to the cluster color in the UMAP. The edge represents the similarity between terms and shorter and thicker edge indicates higher similarity. The full list of GO terms of all the clusters is available in Supplementary Table S3. Scale bars, 20 µm (**a**, **b**, **e**).

**Supplementary Figure 7.**
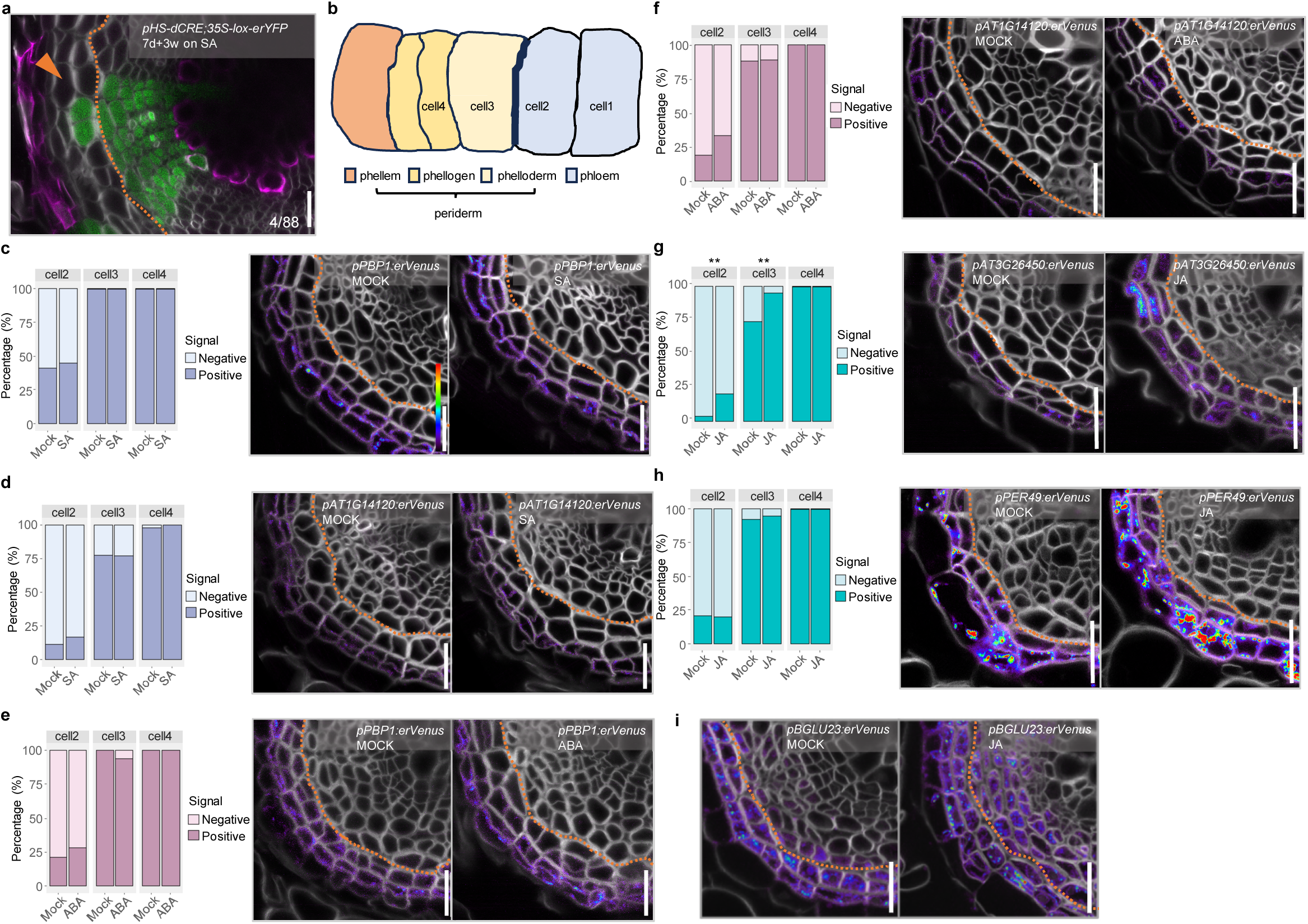
Obtaining periderm identity upon stress hormone treatment. **a**, Cross-section of *pHS-dCRE*;*35S-lox-erYFP* root. Clones induced in 7-day-old seedlings were analyzed 3 weeks after growth on 1/2 GM plates supplemented with 10 µM SA. The orange arrow indicates the transition sector. The fraction indicates 4 out of 88 sectors extended to the phelloderm. Lignin stained with basic fuchsin were shown in magenta**. b**, Schematic illustration of cell-number definition in a cell file for quantification. Cell1 and 2 are mature phloem parenchyma cells and cell3 and 4 are periderm cells. Cell2 is the most distal mature phloem parenchyma cell next to the periderm and cell 1 is the parenchyma cell proximally adjacent to cell2. Cell3 is the most proximal phelloderm cells next to the mature phloem parenchyma and cell 4 is the periderm cell distally adjacent to cell 3. **c**, **d**, Confocal cross-sections of 12-day-old *pPBP1:erVenus* (**c**) and *pAT1G14120:erVenus* (**d**) root grown without (MOCK) or with (SA) 10 µM SA for 2 days and the ratio of signal positive cells in each numbered-cell defined in **b**. **e**, **f**, Confocal cross-sections of 12-day-old *pPBP1:erVenus* (**e**) and *pAT1G14120:erVenus* (**f**) root grown without (MOCK) or with (ABA) 10 µM ABA for 2 days and the ratio of signal positive cells in each numbered-cell defined in **b**. **g**, Confocal cross-sections of 12-day-old *pAT3G26450:erVenus* (**g**) and *pPER49:erVenus* (**h**) roots grown without (MOCK) or with (JA) 10 µM JA for 2 days and the ratio of signal positive cells in each numbered-cell defined in **b. i**, Confocal cross-sections of 12-day-old *pBGLU23:erVenus* roots grown without (MOCK) or with (JA) 10 µM JA for 2 days. In **a**, **c**-**i**, orange dashed line indicates the clonal boundary of vascular tissue and periderm. In **c**-**h**, Fisher’s test was performed. \*\**p*<0.01. In cross sections of **c**-**i**, cell walls stained with SR2200 were shown in grey. Scale bars, 20 µm (**c**-**i**).

**Supplementary Figure 8.**
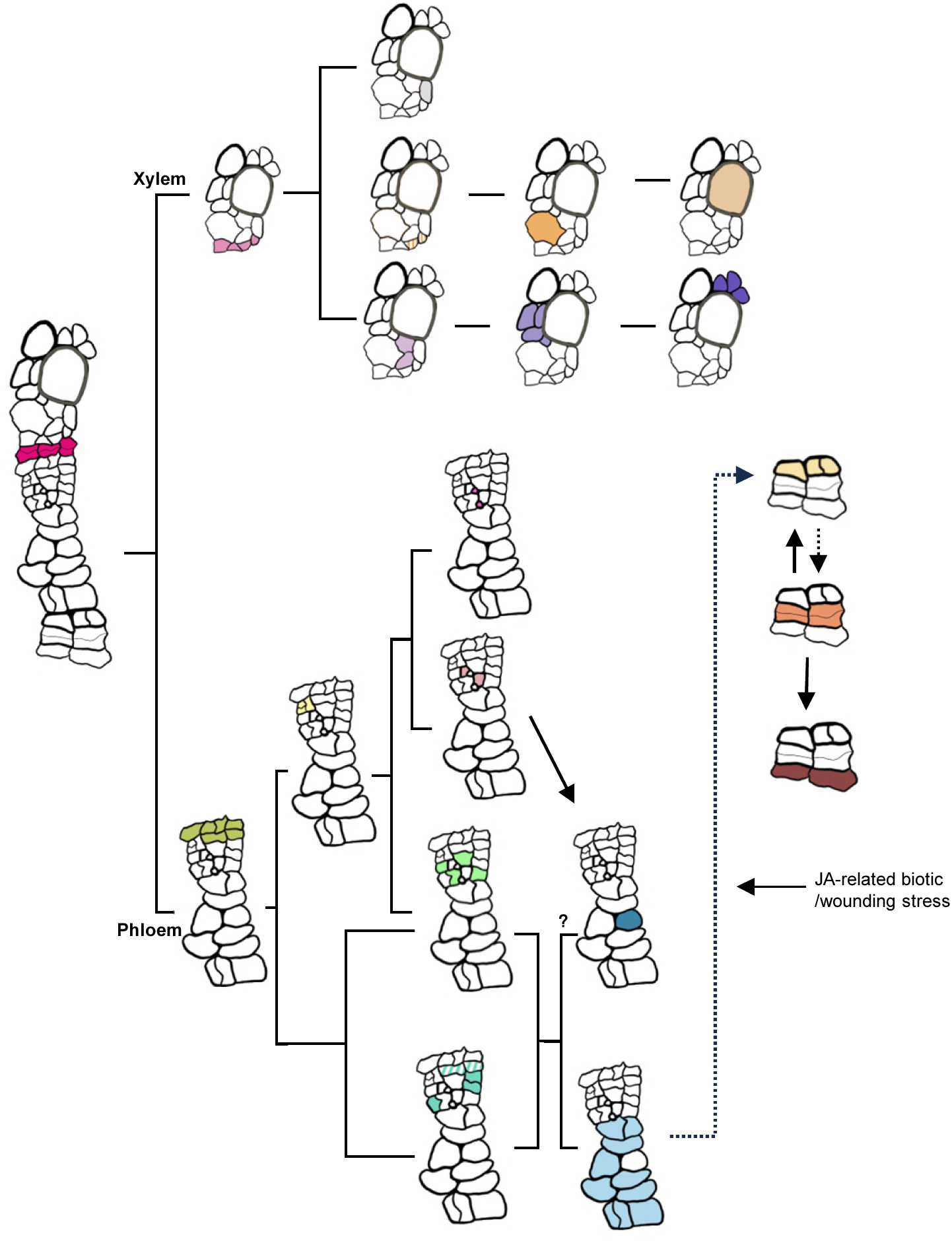
A detailed hierarchical cell fate determination map of the *Arabidopsis* root undergoing secondary growth. Detailed schematic illustration of hierarchical cell fate determination in root secondary tissue. Color of the cell represents cell identity in Fig. 4a.

## Notes

### Competing Interest Statement

The authors have declared no competing interest.

