## Supplementary Notes for "The dynamic and diverse nature of parenchyma cells in the Arabidopsis root during secondary growth"

**Identification of vascular cambium and periderm cells in the scRNA-seq data**

First, we aimed to identify the clusters representing the vascular cambium and cork cambium. We reasoned that these cell types are presented in clusters dominated by cell cycle genes. To investigate this, we initially analyzed the expression of core cell cycle genes^1^ in our dataset. Cluster 15 and 18 were enriched in cells in the G2/M phase, and cluster 13 enriched in cells in the S phase. Because these cell-cycle gene expression were also detected in part of clusters 2 and 6 close to clusters 13 and 15, these cells would also be undergoing cell divisions, forming a ‘meristematic cell band’ in the middle of the UMAP (Extended Data Fig. 1b). Gene Ontology (GO) comparison among differentially expressed genes (DEGs) of clusters 2, 6, 13, 15 and 18 showed cells in cluster 6, 13, 15 and 18 were involved in cell division and pattern specification, while cells in cluster 2 were involved in chemical and hormonal response (Extended Data Fig. 2), suggesting meristematic cells mainly exist in cluster 6, 13, 15, 18 and partially in cluster 2. We selected genes preferentially expressed in the meristematic cell band and examined their expression by generating transcriptional reporter lines. Consistently, promoter activities of *AT2G47780*, *AT2G13820*, *AT1G12080* and *THAUMATIN-LIKE* *PROTEIN-1* (*ATLP-1*) were specifically and *RECEPTOR* *FOR* *ACTIVATED* *C* *KINASE* *1C* (*RACK1C*), *INDOLE-3-ACETIC* *ACID* *12* (*IAA12*), *CELLULASE* *3* (*CEL3*) and *FANTASTIC* *FOUR* *3* (*FAF3*) were preferentially detected in the vascular cambium (Fig. 1c and Extended Data Fig. 1d). Thus, the meristematic cell band clusters (6, 13, 15, 18 and subset of 2) contain the vascular cambium.

Periderm gene, *PEROXIDASE* *15* (*PER15*)^2^ is expressed in cluster 3 and 12, and a small group of dividing cells in clusters 13 and 18 (Extended Data Fig. 1a), genes with similar expression pattern (*PYK10*-*BINDING* *PROTEIN1* (*PBP1*), *PEROXIDASE* *49* (*PER49)*, *BETA*-*GLUCOSIDASE* *23* (*BGLU23*), *AT3G26450*, *AT1G14120* and *VEGETATIVE* *STORAGE* *PROTEIN* *1* (*VSP1*)) showed the expression of reporter gene in the periderm with different tissue preferences (Fig. 1e and Extended Data Fig. 1e). Fluorescence signals in *PBP1* and *BGLU23* reporter lines were detected in the entire periderm; *AT3G26450* and *VSP1* reporter lines showed the highest fluorescence signal intensities in the periderm cells with recently formed cell wall; reporter lines of *PER49* and *AT1G14120* were preferentially expressed in the phellem and phellogen and *PER49* expression peaked in the phellem (Fig. 1e and Extended Data Fig. 1e). The periderm cell types were not separated well between different clusters. *PER49*, *PER15* and *AT1G14120* predominantly expressed in the phellem tended to be detected in cells at the lower side of cluster 3 and 12, suggesting that the region rather than either cluster represents the phellem identity. The phellogen (cork cambium) and phelloderm identities were not clearly distinguishable in the dataset, indicating high similarity between these two periderm cell types. Cluster 4 was close to the periderm clusters, while it shares similar DEGs with cluster 20. Cluster 4- and 20-specific reporters not only showed expression in the periderm, but also in vascular cells (Extended Data Fig. 1f). GO analysis indicated these clusters might consist of cells sensitive to temporal environmental change (Extended Data Fig. 1g).

**New candidate promoters for overexpression analysis in Arabidopsis mature root**

Next, we aimed to identify genes that are broadly and strongly expressed in the secondary tissue. We selected two genes with high number of reads in bulk RNA-seq data^3^ and broad and strong expression in the scRNA-seq data of the mature root. *RIBOSOMAL* *PROTEIN* *S8A* (*RPS8A*) and *UBIQUITIN* *11* (*UBQ11*) promoters drove *YFP* expression strongly and more ubiquitously in the secondary tissue than the commonly used promoter for overexpression, cauliflower mosaic virus 35S (*35S*) (Extended Data Fig. 1h,i). Thus, these two promoters are useful for overexpression studies in root secondary tissues.

**Xylem vessel maturation from committing to terminally differentiating cell and identification of fiber cells**

Based on the expression of previously reported xylem-expressed genes (*NAC*-*DOMAIN* *PROTEIN* *101* (*NAC101*)/*VASCULAR*-*RELATED* *NAC*-*DOMAIN* *6* (*VND6*), *ATHB8*, *NAC*-*DOMAIN* *PROTEIN* *043* (*NAC043*)/*NAC* *SECONDARY* *WALL* *THICKENING* *PROMOTING* *FACTOR1* (*NST1*))^4–6^ (Extended Data Fig. 1a), the clusters above the meristematic cell band (5, 8, 11, 14, 16 and 17) were annotated as xylem clusters. In the dataset, the putative vessel cluster 17 was divided into two subclusters 17_1 and 17_2 (Fig. 1b). GO analysis of the differentially expressed genes (DEGs) revealed that cells in subcluster 17_1 were primarily involved in ribosome biosynthesis, while cells in subcluster 17_2 were undergoing intensive morphological modification, including secondary cell wall biogenesis (Extended Data Fig. 3a). These different GO terms support the idea that there are two stages of vessel differentiation in cluster 17. To further explore the identity of cluster 17, we generated transcriptional fluorescence reporter lines of genes enriched in cluster 17_1, *AT3G10080*, *LIKE* *AUXIN* *RESISTANT* *2* (*LAX2*), *ACAULIS* *5* (*ACL5*), *LACCASE* *6* (*LAC6*), *PHY* *RAPIDLY* *REGULATED* *2* (*PAR2*), *INDOLE*-*3*-*ACETIC* *ACID* *6* (*IAA6*) and *PACLOBUTRAZOL* *RESISTANCE* *5* (*PRE5*). Fluorescence signals in *AT3G10080*, *LAX2*, *ACL5*, and *LAC6* reporter lines were detected in expanding vessel cells and frequently in the cells on the xylem side of the cambium, which had not yet expanded (Extended Data Fig. 3b). In contrast, promoter activities of *PAR2*, *IAA6*, and *PRE5* were detected in the expanding vessels, but not in the xylem side of the cambium (Extended Data Fig. 3b). Because genes detected in the right half of subcluster 17_1 (such as *AT3G10080*, *LAX2*, and *ACL5*) tended to show the expression in the xylem-side cambium in their reporter lines, the small proportion of subcluster 17_1 would represent the earliest stage during commitment to vessel differentiation. Transcriptional reporter lines of genes enriched in subcluster 17_2 (*RHO* *OF* *PLANTS* *GUANINE* *NUCLEOTIDE* *EXCHANGE* *FACTOR* (*ROPGEF4*) and *AT4G27435*) showed strong expression in fully expanded vessels with bright cell wall staining (Extended Data Fig. 3c). In summary, cluster 17 is composed of vessels at different developmental stages; subcluster 17_1 represents vessel identity cells and expanding vessels, followed by differentiating vessels in subcluster 17_2.

Secondary xylem formation in *Arabidopsis* root and hypocotyl occurs in two phases. Initially, vascular cambium produces vessels and parenchyma cells, and when bolting is initiated, fiber cell and vessel formation begins ^7^. Since the seedlings that we used for the scRNA-seq analysis had just initiated bolting, first fiber differentiation events were visible in their roots (Fig. 1a), and xylem fiber regulator *NAC043*/*NST1*^6^ showed high expression in clusters 14 and 16 (Extended Data Fig. 1a). Additionally, the promoter activity of cluster 14- and 16-enriched gene, *CLAVATA3*/*EMBRYO* *SURROUNDING* *REGIONRELATED* *46* (*CLE46)*, was highly detected in differentiating xylem fiber cells (Extended Data Fig. 4a), further supporting the idea that xylem fiber cells are enriched in clusters 14 and 16.

**Secondary sieve element and companion cell maturation resemble the process in primary development**

In the primary root, the phloem tissue consists of conductive sieve elements and companion cells^8^, and *DOF2*.*4*/*PEAR1* is a key regulator and marker in conductive phloem specification^9^. Since *PEAR1* was reported to be expressed in the phloem-side of the cambium^5^, we examined *PEAR1* in our dataset. *PEAR1* was detected in the lower part of the meristematic cell band, preferentially cluster 15, and cluster 21; its transcriptional reporter line showed the highest expression in the phloem-side cambium and the weaker signals in the phloem region along with the conductive phloem formation, suggesting that the *PEAR1*-expressing clusters include phloem identity cells and subsequent daughter cells (Extended Data Fig. 5d). To annotate the conductive phloem clusters in more detail, we investigated the genes that act as downstream of *PEAR1* in the primary phloem^10^. In primary phloem development, *PEAR1* promotes the bifurcation of the procambium and the sieve element lineage by activating *ROPGEF* and *ALTERED* *PHLOEM* *DEVELOPMENT* (*APL*) expression, respectively^10^. We found that *ROPGEF2* and *ROPGEF3* showed specific expression in a subgroup of cells in cluster 15 (Extended Data Fig. 5a). The transcriptional reporter lines of these genes showed strong fluorescence signals in secondary phloem cells undergoing formative division to form sieve elements and companion cells (Extended Data Fig. 5a). These results imply common factors regulating asymmetric cell divisions in primary and secondary phloem formation. *APL* and *SUCROSE*-*PROTON* *SYMPORTER* *2* (*SUC2*), known conductive-phloem marker genes^11,12^, were highly detected in cluster 21 and 19 (Extended Data Fig. 1a). Reporter analysis of genes specific to cluster 21 (*SYTD*, *MAPKK* *KINASE* *20* (*MAPKKK20*) and *AT5G48060*) or cluster 19 (*ETHYLENE*-*RESPONSIVE* *ELEMENT* *BINDING* *FACTOR* *14* (*ERF14*), *AT2G48090*^13^, and *ISOPENTENYLTRANSFERASE* *3* (*IPT3*)^14^) confirmed that cluster 21 and 19 consist of sieve elements and companion cells, respectively (Extended Data Fig. 5b, c).
