## Supplementary Table S1 for "The dynamic and diverse nature of parenchyma cells in the Arabidopsis root during secondary growth"

**Supplementary Table S1: Necessary reported information to allow evaluation and repetition of a plant single-cell/nucleus experiment**

|  | **Details** | **Experimental information** |
| --- | --- | --- |
|  |  | **Secondary root growth** |
| **Biological material** | Species | *Arabidopsis thaliana* |
|  | Accession | Col-0 |
|  | Genotype | WT |
|  | Tissue type | Secondary root |
|  | Detailed growth conditions | Details described in materials & methods |
|  | Harvest conditions | Room temperature |
| **Sample preparation** | Isolation protocol | Details described in materials & methods |
|  | Tissue dissection | Details described in materials & methods |
|  | Fixation | - |
|  | Cell/nuclei enrichment | FACS (BD Aria II) |
|  | Total sample preparation time |  |
|  | Estimated cell/nuclei number loaded | 16000 cells |
|  | Instrument/Method/Kit | Chromium Next GEM Single Cell 3'Kit v3 |
|  | Cell viability test | - |
| **Libraries** | Library construction | According to manufacturer’s instructions. 11 cycles were used for cDNA amplification and 12 for index PCR |
|  | Amplification method | - |
|  | End bias | 3’ |
| **Sequence results** | Instrument/method | Illumina HiSeq 4000 |
|  | Library layout/paired-end | Single index |
|  | N° sequenced reads | 577,256,270 |
| **Raw data** | Reference genome | TAIR10 |
|  | Annotation version | release 40 |
|  | Mapping method (incl. software, customized settings) | Cellranger 6.1.2 |
|  | Mapping efficiency | 87.9% |
|  | Sequencing saturation | 46.1% |
|  | Estimation of ambient RNA | - |
|  | Imputation method and settings | - |
| **Processed data** | N° captured cells | 17,140 |
|  | N° high quality cells | 11,760 |
|  | Filter criteria: % mitochondrial reads/cell or nucleus | Mitochondrial % < 15% |
|  | Filter criteria: % chloroplast reads/cell or nucleus | Chloroplast % < 15% |
|  | Filter criteria: Minimum N° UMI/cell or nucleus | 950 < genes < 8500  4000 < UMIs < 84000 |
|  | N° total detected transcripts | 25,048 |
|  | Doublet rate | - |
|  | Replicate comparisons | - |
|  | Batch correction method for merging (incl. reasoning for batch correction) | - |
|  | Additional processing | - |
| **Validation** | Method of automatic annotation of clusters | - |
|  | Method of manual annotation (markers, gene function info) | Marker lines |
|  | Verification in planta (e.g. Number of markers used for validation) | 93 marker lines |
| **Data availability** | Analysis scripts & codes (GitHub) |  |
|  | Excel Tables DEG for each cluster |  |
|  | Objects/count matrix in repository (which one, where?) | NCBI GEO GSE270140 |
|  | On-line tool/browser URL | www.single-cell.be/plants |
|  | Cell-level metadata table |  |
| **Additional** | additional comments from the authors |  |
